# Immotile cilia of the mouse node sense a fluid flow–induced mechanical force for left-right symmetry breaking

**DOI:** 10.1101/2022.04.11.487968

**Authors:** Takanobu A. Katoh, Toshihiro Omori, Katsutoshi Mizuno, Xiaorei Sai, Katsura Minegishi, Yayoi Ikawa, Hiromi Nishimura, Takeshi Itabashi, Eriko Kajikawa, Sylvain Hiver, Atsuko H. Iwane, Takuji Ishikawa, Yasushi Okada, Takayuki Nishizaka, Hiroshi Hamada

**Author notes:** Correspondence: T.A.K., T.O., or H.H.

## Abstract

Immotile cilia of crown cells at the node of mouse embryos are required for sensing of a leftward fluid flow^1^ that gives rise to the breaking of left-right (L-R) symmetry^2^. The flow-sensing mechanism has long remained elusive, however, with both mechanosensing and chemosensing models having been proposed^1, 3–5^. Here we show that immotile cilia at the mouse node respond to mechanical force. In the presence of a leftward flow, immotile cilia on the left side of the node bend toward the ventral side whereas those on the right side bend toward the dorsal side. Application of mechanical stimuli to immotile cilia along the dorsoventral axis by optical tweezers induced Ca^2+^ transients and degradation of *Dand5* mRNA—the first known L-R asymmetric molecular events—in the targeted cells. The Pkd2 channel protein was found to be preferentially localized to the dorsal side of immotile cilia on both left and right sides of the node, and the observed induction of Ca^2+^ transients preferentially by mechanical stimuli directed toward the ventral side could explain the differential response of immotile cilia to the directional flow. Our results thus suggest that immotile cilia at the node sense the direction of fluid flow in a manner dependent on a flow-generated mechanical force.

---

Breaking of left-right (L-R) symmetry depends on a unidirectional fluid flow at the L-R organizer in fish, amphibians, and mammals^6, 7^, but not in reptiles and birds^8, 9^. In the mouse embryo, the leftward flow at the ventral node (the L-R organizer in this species) is generated by clockwise rotation of motile cilia on pit cells located in the central region of the node. This unidirectional flow is sensed by immotile (primary) cilia on crown cells located at the periphery of the node^2^. How the embryo senses this fluid flow and why only the left-side cilia can respond have remained unknown. Although mechanosensing and chemosensing have each been proposed to underlie this process^3^, the precise mechanism has remained elusive largely as a result of technical difficulties.

## Immotile cilia at the node of mouse embryos undergo asymmetric deformation along the D-V axis in response to nodal flow

We first examined how immotile cilia behave in response to the nodal flow *in vivo*. High-speed live fluorescence imaging revealed that nodal flow induces frequent small bending movements of both left- and right-side cilia (Extended Data Fig. 1, Supplementary Video 1), suggesting that continuous time-independent motion, rather than bilaterally equal periodic motion, contributes to L-R symmetry breaking. To examine whether immotile cilia undergo steady-state deformation in response to nodal flow, we compared the shape of the same cilium in the presence or absence of the flow. Immotile cilia at the node were labeled with mNeonGreen, whereas the cytoplasm of crown cells was labeled with tdKatushuka2 (Fig. 1a). Motile cilia at the node were immobilized by ultraviolet (UV) irradiation, which is thought to induce cleavage of dynein heavy chains^10^, and the shape of immotile cilia was observed by high-resolution microscopy before and after such irradiation (Fig. 1b, c; Extended Data Fig. 2). Nodal flow, as revealed by particle image velocimetry (PIV) analysis, was completely lost after UV irradiation for 45 s (Extended Data Fig. 2a, b; Supplementary Video 2). The flow-dependent bending angle of an immotile cilium was then estimated by ellipsoidal fitting of the shape of the same cilium before and after UV irradiation (Fig. 1d–f, Supplementary Video 3). Examination of the bending angle of immotile cilia of embryos at the two-somite stage revealed that cilia on the left side bent toward the ventral side by 5.0° ± 9.2° (mean ± s.d., *n* = 21), whereas those on the right side bent toward the dorsal side by 4.2° ± 7.4° (*n* = 18) (Fig. 1g, h). This difference in bending angle depended on the presence of the leftward flow, given that it was lost in *iv/iv* mutant embryos (Fig. 1h), which lack nodal flow, and it was dependent on embryonic stage (Fig. 1h). The bending angle was thus significantly asymmetric along the dorsoventral (D-V) axis at the two- and three-somite stages, when the velocity of nodal flow is maximal, whereas it did not manifest asymmetry at the late headfold (LHF) and zero-somite stages, when the flow is absent or weak, respectively^11^ (Fig. 1h). This asymmetric bending of immotile cilia along the D-V axis is consistent with the direction of the flow. Indeed, there was no significant asymmetry in the bending angle along the anterior-posterior (A-P) axis of embryos examined between the LHF and three-somite stages (Fig. 1i, Extended Data Fig. 2f). Of note, immotile cilia with the largest extent of ventral bending were preferentially observed at the left-posterior region of the node (Fig. 1g, Extended Data Fig. 2e), where the first molecular asymmetry appears^12^.

**Fig. 1.**
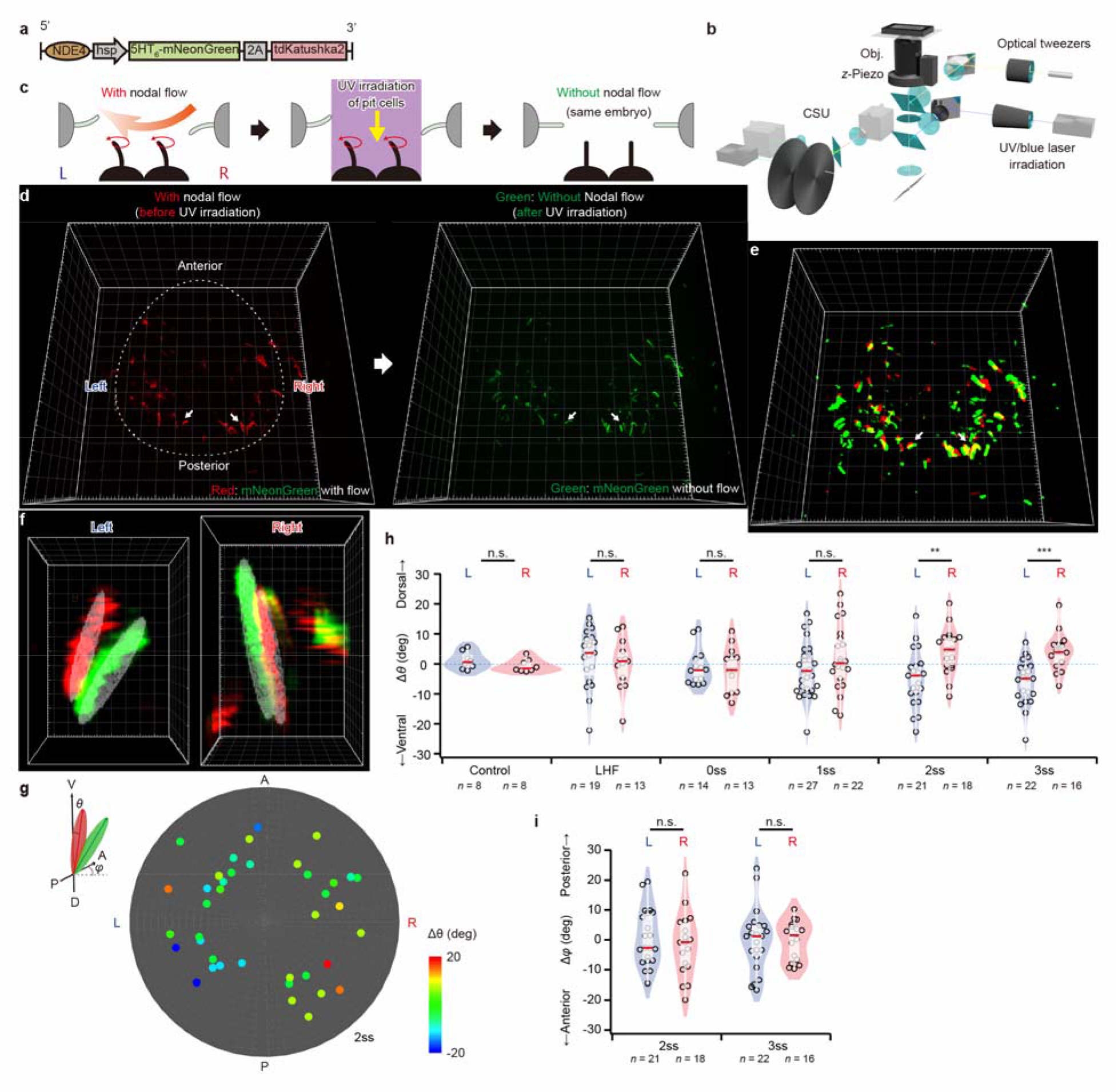
Immotile cilia at the node of mouse embryos undergo asymmetric deformation along the D-V axis in response to nodal flow. **a**, Schematic of the transgene adopted for imaging of immotile cilia. Immotile cilia at the node are visualized on the basis of mNeonGreen expression that is under the control of the crown cell–specific enhancer of the mouse *Nodal* gene (NDE) and is targeted to cilia by a 5-hydroxytryptamine receptor isoform 6 (5HT_6_) sequence. **b**, Schematic of the optical pathway for analysis. Green and red fluorescence are observed simultaneously by spinning-disk confocal microscopy (CSU). A UV laser for irradiation and blue laser for whole-cell fluorescence recovery after photobleaching (FRAP) analysis (see Fig. 2) are introduced into the microscope with a beam expander and iris to control the irradiation area. The microscope is also equipped with optical tweezers and a piezo actuator for mechanical manipulation. Obj., objective lens. **c**, Schematic of the experiment. Live fluorescence images of immotile cilia at the node were first obtained in the presence of nodal flow. The central region of the node encompassing pit cells was then subjected to UV irradiation to abolish nodal flow, and fluorescence images of immotile cilia in the absence of the flow were obtained from the same embryo. **d**, High-resolution three-dimensional (3D) images obtained by deconvolution processing of immotile cilia in the presence (left) or absence (right) of nodal flow. Grid size, 10 µm. **e**, Detection of the edge of each cilium after alignment. Immotile cilia of the same embryo are shown in the presence (red) and absence (green) of the flow. Grid size, 10 µm. **f**, Individual immotile cilia on the left or right side of the node in the presence (red) or absence (green) of the flow. The zenith and azimuth angles were determined by ellipsoidal fitting (gray mesh) of the edge of each cilium. Grid size, 1 µm. **g**, Distribution of immotile cilia at the node with various values of Δθ (change in the zenith angle in response to the flow). Immotile cilia with a lower Δθ value for bending toward the ventral side in response to the flow are preferentially located at the left-posterior region of the node. Data were obtained from wild-type embryos at the two-somite (2ss) stage (see also Extended Data Fig. 2e) (*n* = 39 cilia from seven embryos). **h**, Δθ values (deg, degrees) for immotile cilia on the left and right sides of the node at various developmental stages. Data for *iv/iv* embryos at the LHF to 2ss are shown as a control. Red bars indicate median of samples. **i**, Δφ (change in the azimuth angle along the A-P axis in response to the flow) values for immotile cilia on the left and right sides of the node at the two-and three-somite stages (see also Extended Data Fig. 2f). \*\**P* < 0.01, \*\*\**P* < 0.001; n.s., not significant (Mann-Whitney *U* test).

## Mechanical stimuli administered to immotile cilia by optical tweezers trigger *Dand5* mRNA degradation and Ca^2+^ transients

Given that immotile cilia on the right and left sides of the node were found to bend asymmetrically along the D-V axis in response to the leftward fluid flow, we next tested whether immotile cilia at the node respond to mechanical force with the use of optical tweezers ^13, 14^(Fig. 1b, Extended Data Fig. 3). We examined mouse embryos harboring two transgenes (Fig. 2a, b; Supplementary Video 4): one to visualize perinodal immotile cilia, and the other to monitor the response to mechanical stimuli. *Dand5* mRNA is the ultimate target of nodal flow^15, 16^, being degraded by the Bicc1-Ccr4 complex in response to the flow^17^. Crown cells were labeled with the *NDE4-hsp-dsVenus-Dand5-3′-UTR* transgene, with the level of dsVenus mRNA reflecting that of *Dand5* mRNA^17^, whereas their immotile cilia were visualized with a transgene encoding mCherry. We applied whole-cell fluorescence recovery after photobleaching (FRAP) to examine the kinetics of *Dand5* mRNA after subjecting an immotile cilium to mechanical stimuli (Extended Data Fig. 4a). We first tested the validity of the whole-cell FRAP system. According to a theoretical model, the rate of fluorescence recovery depends on the mRNA level (Extended Data Fig. 4b). In wild-type embryos with a leftward flow, the level of fluorescence on the right side of the node rapidly recovered after photobleaching, consistent with the predicted FRAP curve, whereas the extent of recovery was much lower on the left side (Extended Data Fig. 4c). These observations confirmed that the whole-cell FRAP system was indeed able to monitor the kinetics of *Dand5* mRNA.

**Fig. 2.**
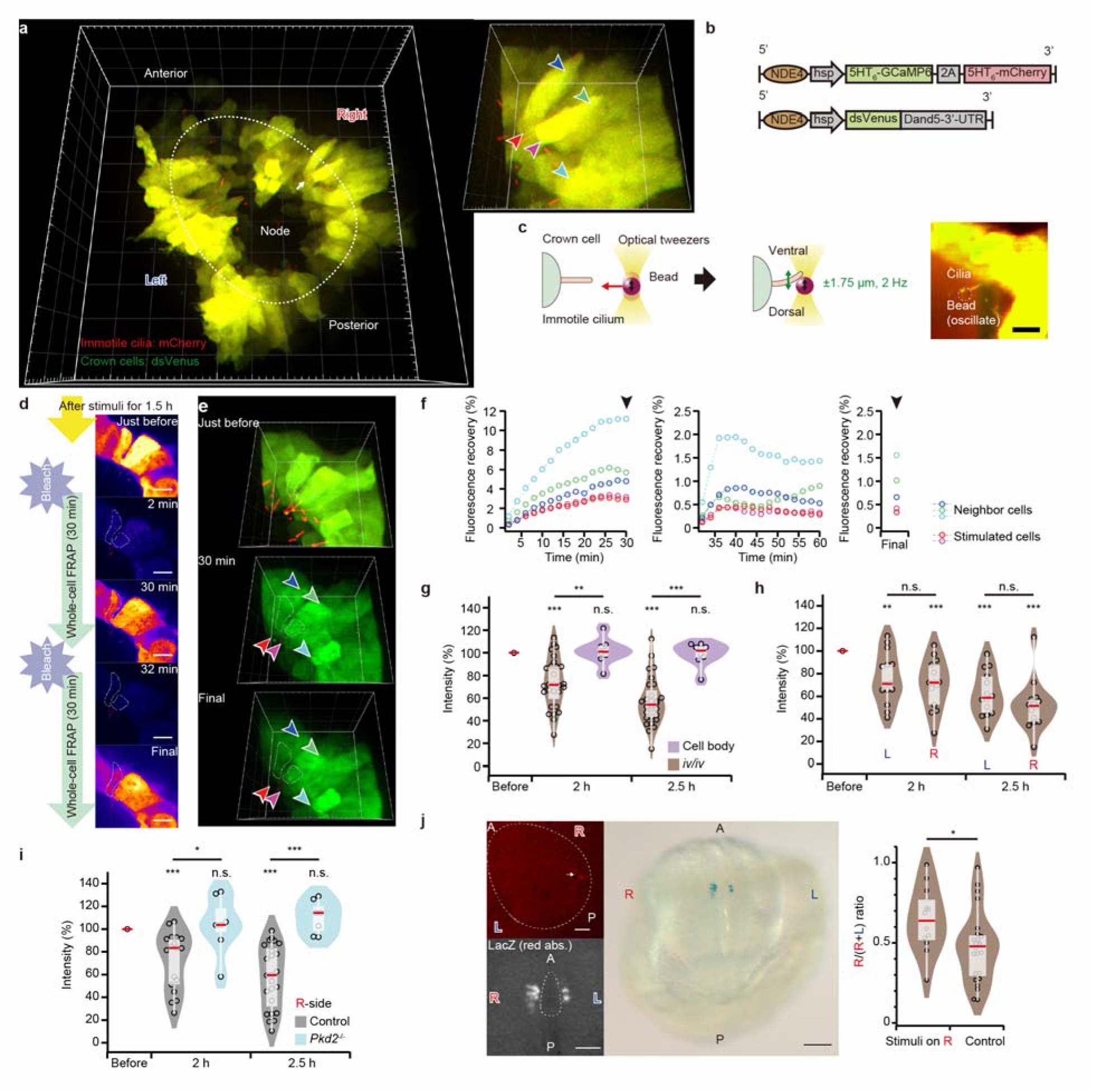
Mechanical stimuli administered to immotile cilia by optical tweezers trigger *Dand5* mRNA degradation and increase Nodal activity. **a**, A 3D image of the node of an *iv/iv* mouse embryo at the two-somite stage harboring the two transgenes shown in **b**. mCherry (red) marks immotile cilia, whereas *Dand5* mRNA degradation in crown cells can be monitored by measurement of dsVenus fluorescence (green). Grid size, 20 μm. The cells to which mechanical stimuli were applied (red and purple arrowheads) and surrounding unstimulated cells (blue, green, and cyan arrowheads) are shown at higher magnification to the right of the main image. Grid size, 10 μm. **b**, Schematic of the two transgenes adopted for these experiments. The expression of dsVenus in crown cells is regulated by the 3′ untranslated region (UTR) of *Dand5* mRNA, whereas GCaMP6 and mCherry are expressed in immotile cilia of these cells. **c**, Experimental scheme. A polystyrene bead (diameter of 3.5 µm) is trapped, placed into contact with an immotile cilium, and forced to oscillate along the D-V axis at 2 Hz and with an amplitude of ±1.75 µm for 1.5 h with the use of optical tweezers. The right image shows that an oscillating bead (white dotted line) makes contact with cilia (see Supplementary Video 5). **d**–**h**, Analysis of *iv/iv e*mbryos (which lack nodal flow) at the EHF to three-somite stage. **d**, *Dand5* mRNA degradation was monitored by whole-cell FRAP (see Extended Data Fig. 4). The entire area of the targeted cells was bleached twice with a 30-min interval between sessions, and fluorescence recovery was monitored. White dotted lines indicate the stimulated cells. 2D sections obtained from 3D images are shown. Scale bars, 10 μm. **e**, 3D images obtained during FRAP. Red, purple, blue, green, and cyan arrowheads represent the same cells shown in **a**. Grid size, 10 µm. **f**, Time course of fluorescence recovery during the first and second FRAP periods. The fluorescence intensity for each cell was measured with a time resolution of 2 min. A 3D image was obtained with a longer exposure time at the end of the second FRAP session (see also Extended Data Fig. 4d). Normalized intensity was calculated with the use of the values indicated by the closed arrowheads and is shown in **g**. **g**, Normalized fluorescence intensity of dsVenus (ratio of the fluorescence intensity of a stimulated cell to that of two or three neighboring unstimulated cells) for before as well as 2 and 2.5 h after stimulation (brown, *n* = 28 embryos). Data are also shown for cells whose cell body (instead of the cilium) was stimulated (purple, *n* = 8 embryos). **h**, Normalized fluorescence intensity of dsVenus before and after stimulation for immotile cilia on the left and right sides (*n* = 14 embryos each for left-side cilia and right-side cilia). **i**, Normalized fluorescence intensity of dsVenus for similar FRAP experiments performed with *Pkd*2^−/−^ and control (wild-type, *iv/+*, or *Pkd2*^+/–^) embryos (*n* = 7 for *Pkd*2^−/−^ and 22 for control embryos). The experiments were performed only with immotile cilia on the right side in order to avoid the effect of nodal flow. \**P* < 0.05, \*\**P* < 0.01, \*\*\**P* < 0.001; n.s., not significant in **g-i** (Mann-Whitney *U* test for different categories of data; Wilcoxon signed-rank test for comparison of before to 2 h and 2.5 h data). **j**, A single cilium on the right side of an *iv/iv* embryo harboring the *ANE-LacZ* transgene was subjected to mechanical stimulation for 1.5 h (arrow in the mCherry fluorescence image shown in the upper left panel), cultured for ∼7 h, and then subjected to X-gal staining to detect Nodal activity (middle panel). The level of staining was quantified as red absorbance (lower left panel), and the ratio of the staining level on the right side to that on the right plus left sides of the node was determined (right panel) (*n* = 12 for stimulated and 24 for control embryos). Scale bars, 20 µm (upper left), 50 µm (lower left), and 100 µm (middle). \**P* < 0.05 (Mann-Whitney *U* test)

Mechanical stimuli were administered under a condition that mimics nodal flow to individual immotile cilia of *iv/iv* embryos (which lack nodal flow) by positioning a polystyrene bead (diameter of 3.5 µm) trapped by optical tweezers into contact with the cilium and displacing it 1.75 μm toward the ventral side and then 1.75 μm toward the dorsal side at a frequency of 2 Hz (Fig. 2c, Supplementary Video 5). The maximal trapping force of ∼±12 pN was sufficient to apply mechanical bending to an immotile cilium (Extended Data Fig. 3a), and observation of beads by 3D single-particle tracking microscopy^18, 19^ confirmed that they indeed moved along the D-V axis (Extended Data Fig. 3c). The infrared laser of the optical tweezers did not exert any unexpected effects such as a change in Ca^2+^ oscillation pattern in crown cells or cilia (Extended Data Fig. 5, see Methods). After administration of mechanical stimuli to an immotile cilium for 1.5 h, all crown cells were subjected twice to uniform photobleaching with a recovery period of 30 min after each bleaching. The recovery of dsVenus fluorescence in the cell with the stimulated cilium and in neighboring unstimulated cells was monitored by time-lapse 3D imaging (Fig. 2d, e; Supplementary Video 6). The fluorescence intensity at 30 min after each bleaching had reached a plateau and was compared between stimulated and neighboring cells (Fig. 2f, Extended Data Fig. 4d). The extent of fluorescence recovery in the stimulated cell was substantially lower than that in the unstimulated cells, with values of 69.9 ± 21.5% at 2 h and 53.0 ± 20.9% at 2.5 h after the onset of stimulation (geometric means ± s.d., *n* = 28) (Fig. 2g), suggesting that mechanical stimulation of an immotile cilium was able to induce degradation of *Dand5* mRNA. Immotile cilia on the right and left sides of the node of *iv/iv* embryos responded similarly to the mechanical stimuli (Fig. 2h). Those on the right side of the node of wild-type embryos (in the presence of the endogenous nodal flow) also showed a similar response to mechanical stimuli (Fig. 2i, Extended Data Fig. 4e–g, Supplementary Video 7). Administration of mechanical stimuli to the cell body of a crown cell instead of to its cilium did not affect recovery of the fluorescence signal (Fig. 2g). Furthermore, the response to mechanical stimuli was lost in embryos lacking the cation channel Pkd2 (Fig. 2i), suggesting that mechanical stimulation of immotile cilia induces degradation of *Dand5* mRNA in a Pkd2-dependent manner.

The response to mechanical stimuli was further confirmed with another readout, expression of the transgene *ANE-LacZ*, which allows monitoring of Nodal activity in perinodal cells^20^. This transgene manifests left-sided expression at the node in the presence of nodal flow, but shows L-R randomized expression in its absence^12^. Mechanical stimulation along the D-V axis of an immotile cilium on the right side of an *iv/iv* embryo resulted in a significant increase in the R/(R + L) ratio of *ANE-LacZ* expression to 0.64 ± 0.19 (mean ± s.d., *n* = 12), compared with a value of 0.47 ± 0.22 (*n* = 24) for control embryos (Fig. 2j). These results suggested that mechanical stimulation of a single immotile cilium not only induced degradation of *Dand5* mRNA in the targeted cell but also established molecular asymmetry in all crown cells at the node. A feedback mechanism involving Wnt and Dand5 signals^15^ may be responsible for expansion of asymmetric Nodal activity among crown cells.

Perinodal cells of mouse embryos manifest both cytoplasmic and intraciliary Ca^2+^ transients in response to nodal flow^21–23^. We therefore examined whether mechanical stimuli administered to immotile cilia of *iv/iv* embryos might induce such transients. Cytoplasmic and intraciliary Ca^2+^ transients were observed with cytoplasm-targeted and cilium-targeted forms of the fluorescent Ca^2+^ indicator GCaMP6, respectively (Fig. 3a, b). Spontaneous Ca^2+^ transients were detected in both the cytoplasm and immotile cilia (Fig. 3c), with such transients having been shown to be independent of nodal flow and the Pkd2 channel^21, 22^. However, the frequency of Ca^2+^ transients increased significantly from 0.83 ± 0.71 to 1.39 ± 1.65 spikes/min in the cytoplasm (means ± s.d., *n* = 42 cells) and from 0.32 ± 0.57 to 0.59 ± 1.02 spikes/min in cilia (*n* = 24) in response to mechanical stimuli (Fig. 3c, d; Supplementary Video 8). Such increases in the frequency of cytoplasmic and ciliary Ca^2+^ transients were not observed in embryos lacking the Pkd2 channel (Fig. 3e).

**Fig. 3.**
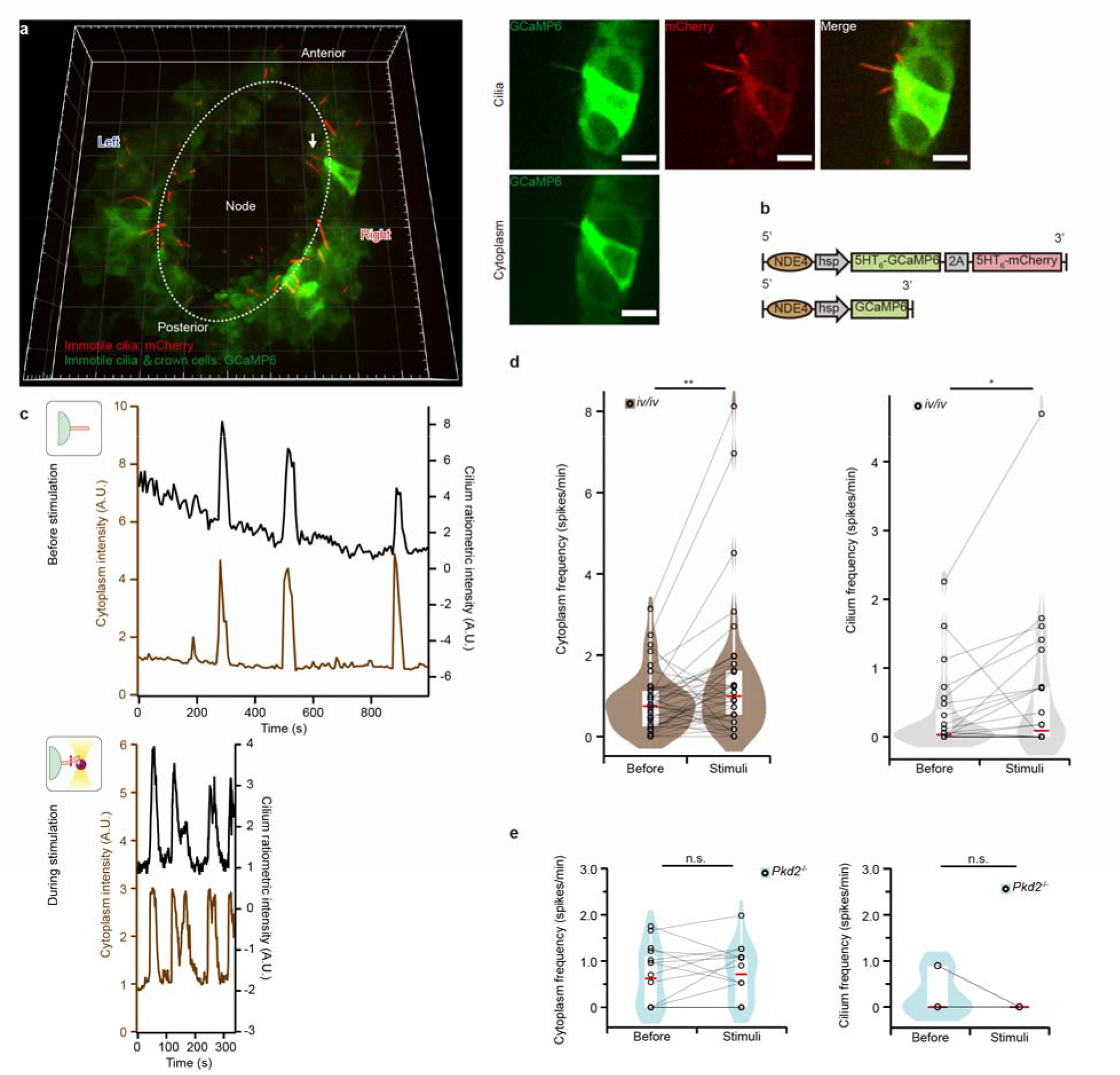
Mechanical stimulation of the immotile cilium of crown cells alters the dynamics of Ca^2+^ signaling in both the cilium and cytoplasm. **a**, A 3D image of the node of an *iv/iv* embryo at the two-somite stage harboring the two transgenes in **b** is shown on the left. Grid size, 10 µm. Both GCaMP6 and mCherry are expressed in immotile cilia for ratiometric Ca^2+^ imaging, with sections containing the cilium (usually three sections containing a cilium located ventrally relative to the cell body) being averaged (upper right panels). GCaMP6 is also expressed in the cytoplasm for cytoplasmic Ca^2+^ imaging, with either a single section being analyzed or, occasionally, sections containing the cell body being averaged (lower right panel). Scale bars, 10 μm. **b**, Schematic of the two transgenes used for intracellular Ca^2+^ measurement. **c**, Time course of Ca^2+^ signal intensity before (upper panel) and during (lower panel) stimulation of the immotile cilium of a crown cell in an *iv/iv* embryo. Brown traces indicate cytoplasmic Ca^2+^ (*F*/*F*_0_ ratiometric values), whereas black traces indicate intraciliary Ca^2+^ (GCaMP6/mCherry *F*/*F*_0_ ratiometric values). Calcium dynamics in the cilium and cytoplasm were monitored for ∼15 min before (upper) and then for ∼5 min after (lower) the onset of mechanical stimulation of the cilium. See Supplementary Video 8. **d**, Mean frequency of Ca^2+^ transients in the cytoplasm (left) and cilium (right) measured as in **c** (*n* = 42 cells from 28 embryos for cytoplasm and 24 cilia from 17 embryos for cilia). \**P* < 0.05, \*\**P* < 0.01 (Wilcoxon signed-rank test). **e**, Mean frequency of Ca^2+^ transients in the cytoplasm and cilium of *Pkd*2^−/−^ embryos (*n* = 16 cells from 16 embryos for cytoplasm and 6 cilia from six embryos for cilia). n.s., Wilcoxon signed-rank test.

## Immotile cilia sense bending direction in a manner dependent on polarized localization of Pkd2

We next asked whether an immotile cilium might respond differentially to forced bending toward the dorsal or ventral sides, possibly as a result of a structure or molecule within the cilium that can sense the direction of bending. The polarized distribution of such a structure or molecule relative to the midline of an embryo would allow a differential response to the direction of nodal flow (Extended Data Fig. 6a). We first searched for such a structure at or near the base of immotile cilia by focused ion beam–scanning electron microscopy (FIB-SEM), which would be expected to reveal an anisotropic distribution relative to the midline, but we were not successful (Extended Data Fig. 6b). Consistent with this result, measurement of the flexural rigidity of immotile cilia with optical tweezers revealed no apparent difference between dorsal and ventral bending (Extended Data Fig. 7h).

Alternatively, a mechanosensitive channel may be preferentially localized to one side (dorsal or ventral) of an immotile cilium. The most likely candidate for such a channel would be Pkd2, given that the ciliary localization of this protein is essential for breaking of L-R symmetry^2, 24^. We examined the precise localization of Pkd2 within immotile cilia by 3D stimulated emission depletion (STED) microscopy, with wild-type embryos harboring an *NDE2-hsp-Pkd2-Venus* transgene that can rescue the defects of Pkd2 deficient mouse embryos^2^. The super-resolution images showed non-uniform distribution of Pkd2∷Venus protein on each immotile cilium. The Pkd2::Venus protein accumulated to form clusters on the surface of the cilium. More importantly, the cluster was preferentially localized to the dorsal side (the side facing the midline of the embryo) of immotile cilia on both the left and right sides of wild-type embryos (Fig. 4a; Supplementary Video 9). Analysis of the angular distribution of Pkd2::Venus protein on the transverse plane of the axoneme revealed that the intensity of the Pkd2 signal on the dorsal side (0.54 ± 0.12) was significantly stronger than that on the ventral side (0.46 ± 0.12) (means ± s.d., *n* = 50) (Fig. 4b). Preferential localization of Pkd2 on the dorsal side of immotile cilia was confirmed by confocal microscope with the Airyscan detector. Analysis of the distance along the *z*-axis between the center of localization of Pkd2 and that of the axoneme by Gaussian fitting revealed that the Pkd2 region was thus displaced toward the dorsal side by 142 ± 92 nm in cilia on the left side (mean ± s.d., *n* = 53) and by 192 ± 203 nm in those on the right side (*n* = 54) (Fig. 4c, Extended Data Fig. 8b,c; Supplementary Video 10), with these distances being compatible with a value of 100 nm for the radius of an axoneme. Finally, images of longitudinal sections in immotile cilia further confirmed the dorsal localization of Pkd2 on immotile cilia (Extended Data Fig. 8e). In contrast, there was no significant enrichment of Pkd2 along the proximodistal axis of a cilium (Extended Data Fig. 8d). Enrichment of Pkd2 at the dorsal side of a cilium could explain how immotile cilia sense the direction of nodal flow (Fig. 4d). Indeed, imposition of mechanical stimuli in a single direction and measurement of cytoplasmic Ca^2+^ transients revealed that immotile cilia showed a significantly greater response to stimuli directed toward the ventral side than to those directed toward the dorsal side (Fig. 4e, Extended Data Fig. 9, Supplementary Video 11).

**Fig. 4.**
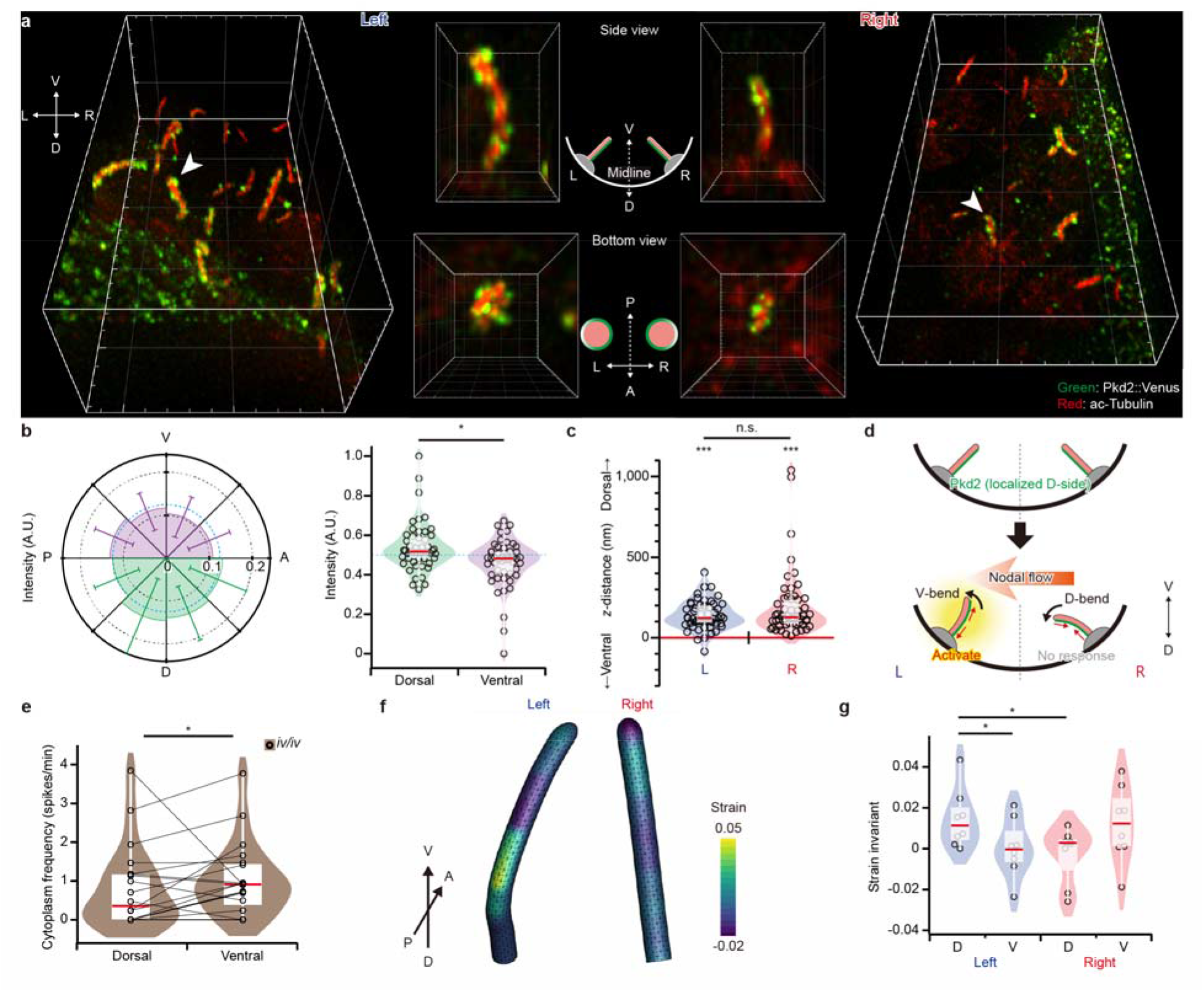
Immotile cilia sense bending direction in a manner dependent on polarized localization of Pkd2. **a**, A wild-type mouse embryo harboring an *NDE2-hsp-Pkd2-Venus* transgene^2^ was subjected to immunofluorescence analysis for detection of the Pkd2::Venus fusion protein and acetylated (ac)–tubulin at the node with an 3D STED microscope (left and right image). Magnified views of D-V sections and bottom views shown in the middle suggested a preferential localization of Pkd2::Venus at the dorsal side of cilia on both the left and right sides of the node. Grid size, 5 µm (main panels), 1 µm (side view), 0.5 µm (bottom view). **b**, Angular distribution of green fluorescence intensity in transverse plane of each cilium imaged as in **a** was analyzed (see Methods). The intensity of Pkd2::Venus on the dorsal side was significantly stronger than that on the ventral side (*n* = 50 cilia from four embryos). \**P* < 0.05 (Wilcoxon signed-rank test). **c,** Distance along the *z*-axis between the centers of red and green fluorescence intensity in longitudinal optical-section of each cilium obtained by confocal microscope with Airyscan detector (see Extended Data Fig. 8b) was measured by Gaussian fitting, after precise correction for chromatic aberration. The intensity center for Pkd2::Venus was significantly polarized toward the dorsal side of cilia on both the left and right sides (*n* = 53 cilia for left side and 54 cilia for right side from 13 embryos). \*\*\**P* < 0.001 (One-sample *t*-test) and n.s. (Wilcoxon signed-rank test for comparison of left and right sides). **d**, A model that would explain why immotile cilia on the left side, but not those on the right side, respond to the leftward fluid flow. Pkd2 is localized at the dorsal side of cilia on both left and right sides of the node. Given that membrane tension on the dorsal side is increased by ventral bending, immotile cilia are able to sense only ventral bending. Nodal flow induces ventral bending only to cilia on the left side, allowing these cilia to be activated by the leftward flow. An artificial rightward flow would activate cilia on the right side. **e**, Frequency of cytoplasmic Ca^2+^ transients in individual immotile cilia subjected to both ventral and dorsal vending (*n* = 18 cilia from 18 *iv/iv* embryos). \**P* < 0.05 (Wilcoxon signed-rank test). **f**, Estimated strain at the membrane of the immotile cilia on the left and right sides of the node shown in Figure 1f. The membrane is modeled as a 2D hyperelastic material, and the contour color indicates the second strain invariant. **g**, Comparison of strain applied to the dorsal and ventral sides of cilia on the left or right sides. The mean value of the second strain invariant was used as the basis for strain measurement. Definition of the dorsal and ventral regions is described in Extended Data Figure 10b. Strain on the dorsal and ventral sides of a left-side cilium is estimated as 0.014 ± 0.013 and 0.000 ± 0.013 (means ± s.d., *n* = 8), respectively, whereas the corresponding values for a right-side cilium are 0.000 ± 0.001 and 0.012 ± 0.017 (*n* = 8). \**P* < 0.05 (Student’s paired *t*-test)

## Discussion

Our results collectively indicate that immotile cilia at the node respond to mechanical force generated by fluid flow. A similar conclusion was obtained with zebrafish embryos in the accompanying paper (Djenoune et al). Given that the relative extent of viscous and bending force is approximately given by length^4^/stiffness^0.25^ (ref. ^25^), the slender shape of a cilium is suited to sensing of a weak flow and transducing the flow signal into strong locoregional strain. Indeed, in the presence of leftward flow, the bending of an immotile cilium on the left side of the node toward the ventral side imposes a strain of 0.014 ± 0.013 (mean ± s.d., *n* = 8) to the dorsal side of the cilium (Fig. 4f, g). The resulting membrane tension, according to a previously described model^26^, would be 1.6 ± 1.6 mN/m (mean ± s.d., *n* = 8) (Extended Data Fig. 10), which may be sufficient to activate dorsally localized Pkd2 and trigger the Ca^2+^ response. In contrast, on the right side of the node, strain at the dorsal side of an immotile cilium is as small as 0.000 ± 0.001 (*n* = 8) and would not support a response (Fig. 4f, g). Our findings thus suggest how immotile cilia sense the direction of nodal flow: Directional information of the flow is geometrically converted to locoregional strain, which is integrated over the polarized area of Pkd2 localization and allows activation only of cilia on the left side, thereby giving rise to robust L-R determination. Several questions remain, the most intriguing of which is how Pkd2 becomes preferentially localized to one side of an immotile cilium. An unknown signal derived from the midline of the embryo may polarize Pkd2 localization. Bone morphogenetic protein (BMP) antagonists expressed at the node are candidates for such a signal, but treatment of embryos with BMP did not affect the localization of Pkd2 at the dorsal side of immotile cilia (Extended Data Fig. 8f). Characterization of the mechanism responsible for the polarized localization of Pkd2 therefore awaits further study.

## Methods

### Mice and transgenes

*Pkd2*^−/−^ and *iv/iv* mice were described previously^27, 28^. *Kif3a*^+/–^ mice (strain B6.129-Kif3a<TM1GSN>/J) were obtained from The Jackson Laboratory. A DNA fragment encoding the 5HT_6_ sequence was kindly provided by T. Inoue (Johns Hopkins University). GCaMP6 was described previously^29^. tdKatushka2, mNeonGreen^30^, and mKeima^31^ constructs were obtained from Addgene. A DNA fragment encoding the 2A sequence (equine rhinitis A virus 2A peptide)^32^ was synthesized as an oligomer. For expression in node crown cells, transgenes were placed under the control of the node-specific enhancer (NDE) of the mouse *Nodal* gene and the *Hsp68* gene promoter^2, 33^. The coding sequence for mNeonGreen was linked to DNA encoding the 5HT_6_ sequence, whereas that for tdKatushka2 was connected to the mNeonGreen sequence via the DNA sequence for the 2A peptide, yielding the *NDE4-hsp-5HT_6_-mNeonGreen-2A-tdKatushka2-pA* transgene. *NDE4-hsp-GCaMP6-pA* and *NDE4-hsp-5HT_6_-GCaMP6-2A-5HT_6_-mKeima-pA* were constructed in a similar manner. For generation of transgenic mice, each transgene was microinjected into the pronucleus of fertilized eggs obtained by crossing C57BL/6J females with C57BL/6J males. Transgenic mouse lines harboring *NDE2-hsp-Kif3a-IRES-LacZ*^2^*, NDE4-hsp-dsVenus-Dand5-3′-UTR*^17^, *NDE4-hsp-5HT_6_-GCaMP6-2A-5HT_6_-mCherry*^21^, *ANE-LacZ*^12^, or *NDE2-hsp-Pkd2-Venus*^2^ were described previously. Expression of *ANE-LacZ* was detected by staining with X-gal (5-bromo-4-chloro-3-indoyl-β-D-galactoside)^33^. All animal experiments were approved by the Institutional Animal Care and Use Committee of RIKEN Kobe Branch.

### Mouse embryo culture

Mouse embryos were collected at embryonic day (E) 7.5 in HEPES-buffered Dulbecco’s modified Eagle’s medium (DMEM, pH 7.2). Those at the early headfold to early somite stages were selected and cultured by the roller culture method under 5% CO_2_ at 37°C in 50-ml tubes containing DMEM or FluoroBrite DMEM (A1896701, ThermoFisher) supplemented with 75% rat serum. Agents were added to the medium at final concentrations of 1 mg/ml for BMP2 (355-BEC-010, R&D Systems), 500 μM for GdCl_3_ (16506-71, Nacalai Tesque), and 1 µM for thapsigargin (T9033, Sigma).

### High-speed imaging of the motion of immotile cilia by HILO microscopy

Highly inclined and laminated optical sheet (HILO) microscopy^34^ was performed with an inverted microscope (1.5× Magnifier, Ti-E; Nikon) equipped with a 60× objective lens (CFI Apo TIRF 60×H 1.49 N.A., Nikon), a 488-nm laser (Sapphire, Coherent) and filter set (LF488/561-A, Semrock), an electron-multiplying charge-coupled device (EM-CCD) camera (2 × 2 Binning mode, iXon^+^ DU897E; Andor) with image-splitting optics (W-VIEW GEMINI, Hamamatsu Photonics), and a stage-top CO_2_ incubator (INUG2-TIZ, Tokai hit). A distal portion of each embryo including the node was excised, placed into a chamber consisting of a glass slide fitted with a thick silicone rubber spacer (thickness of 400 µm) to prevent disturbance of nodal flow, covered with a coverslip (No.1S, Matsunami), and incubated under 5% CO_2_ at 37°C in DMEM supplemented with 75% rat serum. The microscope and camera were controlled by MetaMorph software (Molecular Devices). Sequential images were acquired with a time resolution of 29.2 ms and were analyzed with the use of ImageJ (version 1.52a, NIH) and IgorPro 8 (WaveMetrics) software. Images in the green channel were rotated with bicubic interpolation to align cilia along the *x*-axis. The intensity of 5 pixels from the distal end was averaged, and the location of the ciliary tip along the *y*-axis was tracked by 1D Gaussian fitting (Extended Data Fig. 1d).

### 3D imaging and analysis of flow-dependent passive bending of immotile cilia

Immotile cilia at the node of mouse embryos harboring the *NDE4-hsp-5HT_6_-mNeonGreen-2A-tdKatushka2* transgene were visualized with an inverted microscope (IX83, Olympus) equipped with a spinning-disk confocal unit (CSU-W1, Yokogawa), a 100× lens (UPLAPO100XOHR 1.5 N.A., Olympus), a 375-nm laser (70 mW, JUNO 375; Kyocera SOC) and filter set (zt405/488/561rpc, chroma), a laser beam expander (GBE15-A, Thorlabs), an iris (G061653000, Linos) to allow near-uniform UV laser irradiation of only the node area, a piezo *z*-stage (P-721, Physik Instrumente), and a stage-top CO_2_ incubator (STXF-IX3WX and GM3000, Tokai hit). The node region of the embryo was set on a glass slide as described in the previous section and incubated in FluoroBrite DMEM supplemented with 75% rat serum in order to minimize the background. mNeonGreen was detected with a laser (wavelength of 488 nm) and 520/35 filter set and with an EM-CCD camera (10 MHz A/D was adopted to reduce noise; iXon Ultra 888, Andor) equipped with a water cooler (LTB-125A, ASONE). 3D images were recorded with a depth along the *z*-axis of 200 nm and 100 sections, and the node was then subjected to UV irradiation (23 mW before the objective) for 45 s to stop the beating of motile cilia. 3D images of the same embryo were then acquired again immediately after UV irradiation. The 3D images were analyzed after 3D deconvolution with DeconvolutionLab2^35^ and the Richardson-Lucy algorithm with total variation regularization^36^. The point spread function for deconvolution was obtained with the use of fluorescent beads (diameter of 200 nm; F8811, ThermoFisher). After processing of images by Gaussian filtering, the edge of each cilium was detected and binarized, the *xyz* location of the cilum base was determined and used as a fiducial marker, and the image acquired after UV irradiation was transformed into coordinates. The zenith (θ) and azimuth (φ) angles of each cilium were determined by ellipsoidal fitting, and the change in angle (Δθ and Δφ) was calculated by comparison of the angles before and after UV irradiation of the same cilium. Ellipsoidal fitting of the ciliary shape was performed with the eigenvalues and eigenvectors of the gyration tensor, which is given by

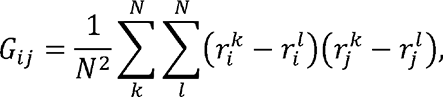

where **r***^k^* is the *k*th material point of the cilium, and the subscript *i* is the vector component in the *i* direction (likewise for *l* and *j*).

### PIV analysis

Nodal flow was observed with an IX83/CSU-W1 microscope equipped with a 60× lens (UPLSAPO60XW 1.2 N.A., Olympus) and EM-CCD camera as described in the previous section, and PIV analysis was performed as described previously^11^. The excised node region was exposed to DMEM supplemented with 75% rat serum and 0.2-µm-diameter fluorescent beads (505/515, F8811; Invitrogen), and the motion of the beads in the node cavity was monitored in planes of ∼10 µm above the surface of node pit cells.

### 3D single-particle tracking microscopy

Validation of movement of a trapped fluorescent bead (diameter of 4 µm; F8858, ThermoFisher) along the *z*-axis (Extended Data Fig. 3) and of force measurement along the *z*-axis (Extended Data Fig. 7) was performed by 3D single-particle tracking microscopy^18, 19^ with a cylindrical lens (*f* = 1000 mm; LJ1516RM-A, Thorlabs) and relay lens (pair of AC254-100-AB, Thorlabs). Calibration was performed with a piezo *z*-stage (P-721, Physik Instrumente), and analysis was performed as described previously^19^ with codes written in IgorPro 8 (WaveMetrics).

### Manipulation of cilia by optical tweezers

The node region of a mouse embryo was set on a glass slide as described above for HILO microscopy and was covered with a thickness-controlled coverslip (No.1S HT, Matsunami). Polystyrene beads (diameter of 3.5 µm; S37224, ThermoFisher) were diluted 1:50 in FluoroBrite DMEM, isolated by centrifugation at 20,300 × *g* for 15 min, and resuspended at a dilution of 1:10. A portion of the diluted beads was transferred to the medium above the node with a P2 pipette tip so as to expose the node to ∼1 to 10 beads. FluoroBrite DMEM supplemented with 75% rat serum was used as the culture medium to reduce absorbance for the infrared laser used for the optical tweezers. Immotile cilia at the node were visualized with an IX83/CSU-W1 microscope equipped with a 60× lens (UPLSAPO 60XW 1.2 N.A., Olympus) and with a single-mode fiber laser (wavelength of 1064 nm; YLR-5-1064-LP-SF, IPG Photonics) and filter set (ZT1064rdc-sp, Chroma Technology, and SIX870, Asahi). A long-path filter was inserted before a halogen lamp (LV0630, Asahi) in order to visualize the bright-field image of beads with the red channel of CSU-W1. A bead was trapped and then oscillated with the piezo *z*-stage (P-721, Physik Instrumente) controlled by LabVIEW (National Instruments) via a digital-analog converter (USB-6363, National Instruments). A trapped bead was manually manipulated by a motorized XY stage (BIOS-225T and FC-101G, Sigma), and contact of the bead with the cilium during oscillation for 1.5 h was monitored by ciliary GCaMP6 or mCherry fluorescence imaging or bright-field imaging of the bead. Calibration of optical tweezers was performed as described previously^37^. For application of stimuli to the cell body, a bead was manipulated so as to make contact with the cell body while avoiding contact with the cilium. For sequential application of stimuli to the dorsal and ventral sides of a cilium, the piezo *z*-stage was moved as shown in Extended Data Figure 9a. For elasticity measurement, a sulfate-modified fluorescent polystyrene bead (F8851, ThermoFisher) was attached to a cilium by exposure of the node to a 1:200 dilution of beads in Ca^2+^-and Mg^2+^-free phosphate-buffered saline for 3 min, and unattached beads were then washed away. The 3D shape of the cilium was measured with the use of CSU-W1, and the optical pathway was then changed to 3D single-particle tracking microscopy as described above. The initial position of the bead attached to the cilium was measured, after which the bead was trapped with a laser poser of 800 mW and moved with the piezo *z*-stage (Extended Data Fig. 7e). Finally, the center of the trapping position was determined by trapping the bead with a laser power of 2 W. Deformation of cilia by optical tweezers was modelled as a cantilevering problem in an Euler–Bernoulli beam. According to the Euler-Bernoulli beam equation, elastic deflection δ at the free-end of a cantilever beam is given by δ = *FL*^3^/3*E*_b_, where *F* is the bending force, *L* is the length of the cilium, *E*_b_ is the bending rigidity. The bending rigidity in the dorsal and ventral directions was estimated by substituting the experimentally obtained deflections and forces into the above equations.

### Analysis of the degradation of *Dand*5 mRNA by whole-cell FRAP

Uniform bleaching of cells at the node was performed with an IX83/CSU-W1 microscope equipped with a 60× lens (UPLSAPO 60XW 1.2 N.A., Olympus), a 488-nm laser (55 mW, Sapphire, Coherent) and filter set (zt405/488/561rpc, Chroma), and a pair of laser beam expanders (LBED-5 and LBED-3, Sigmakoki). Before bleaching, 3D images of ciliary mCherry and cytoplasmic dsVenus were recorded with a *z*-axis depth of 1 µm and 30 sections. For fluorescence bleaching, the node region was irradiated with the 488-nm laser (output power set at 55 mW) for 3 min, after which 3D images of dsVenus were obtained to examine the time course of the fluorescence intensity for 30 min. Bleaching and measurement of fluorescence recovery were then repeated before recording of 3D images with a longer exposure time. The cell with the cilium subjected to mechanical stimuli was carefully tracked in a 3D movie, and the fluorescence intensity of this cell and of surrounding cells was measured in images obtained before stimulation and at 30 min after the first bleaching as well as in the final image. The ratio of intensity of the targeted cell to that of surrounding cells was calculated. The normalized intensity, with the initial ratiometric intensity defined as 100%, is presented. If mRNA is translated to protein with a rate constant γ and protein degradation occurs with a rate constant δ, the relation between the amount of mRNA (*mRNA*) and that of protein (*P*) is described as *P*(t) = –γ/δ [1 – *exp*(–δ*t*)] *mRNA*, where *t* is time (Extended Data Fig. 4b). Given that fluorescence recovery reached a plateau within 30 min (Fig. 2f; Extended Data Fig. 4c, d), the normalized intensity at 30 min (*I*_30_) is linearly related to the initial mRNA level as *I*_30_ ∝ –γ/δ *mRNA*.

### Analysis of Nodal activity

Embryos from the LHF to zero-somite stages were selected and placed on a glass-bottom dish as described previously^38^. Stimulation was performed with optical tweezers as described above, on a glass-bottom dish. After mechanical stimulation of cilia, embryos were collected from the dish and cultured for ∼7 h, also as described above. Embryos were stained with X-gal, and those at the four-to five-somite stage were observed with a stereomicroscope (M205 FA, Leica) equipped with a cooled color CMOS camera (DP74, Olympus). Given that the products of X-gal staining absorb red light, the sum of the decrease in absorbance from baseline for the blue-stained region on the right side and that on the left side was measured with the red channel and the R/(R + L) ratio was estimated.

### Analysis of Ca^2+^ transients

Before application of mechanical stimuli, 3D images of ciliary mCherry and GCaMP6 as well as cytoplasmic GCaMP6 were obtained for 17 min at a time resolution of 6.6 s with an IX83/CSU-W1 microscope equipped with a 60× lens (UPLSAPO 60XW 1.2 N.A., Olympus). During application of the stimuli with optical tweezers as described above, *xy* images of ciliary mCherry and GCaMP6 as well as cytoplasmic GCaMP6 were obtained for 5.5 min at a time resolution of 1.1 s. Note that during administration of stimuli, the piezo *z*-stage is oscillating at 2 Hz with an amplitude of ±1.75 µm and the exposure time of the EM-CCD camera is 1.1 s, which is longer than the period of oscillation; an averaged image in the ±1.75 µm range of the *z*-axis was thus obtained. From the 3D images obtained before stimulation, one or an average of three *z*-stacks containing the cilium or cytoplasm was selected for measurement of intensity. Sections slightly distant ventrally from cells were chosen for measurement of ciliary intensity in order to minimize the leakage of cytoplasmic fluorescence. The intensity of cilia and the cytoplasm during stimulus application was measured from the 2D images acquired during stimulation. The GCaMP6/mCherry ratio and cytoplasmic GCaMP6 intensity were calculated as described previously^21^. *F*_0_ was measured in the resting state and was used to estimate *F*/*F*_0_. The mean frequency denotes the number of spikes per minute.

### Analysis of Pkd2 localization in immotile cilia

E7.5 mouse embryos were recovered in phosphate-buffered saline (PBS), fixed for 1 h at 4°C in PBS containing 4% paraformaldehyde, and permeabilized for 30 min at room temperature in PBS containing 0.2% Triton X-100. The embryos were then incubated overnight at 4°C first with primary antibodies and then with secondary antibodies diluted in PBS containing 0.01% Triton X-100 (PBST). Primary antibodies included those to acetylated tubulin (1:200 dilution; T6793, Sigma) and to green fluorescent protein (1:200 dilution; ab13970, Abcam), and secondary antibodies included anti-chick or anti-mouse conjugated either with AlexaFluor 488 (1:500 dilution; A11039 and A11001, respectively, Invitrogen) or with AlexaFluor 568 (1:500 dilution; A11041 and A11004, respectively, Invitrogen). After incubation with secondary antibodies, the embryos were washed six times (1 h each time) with PBST. For preparation of frozen sections of the node, embryos were transferred to a series of sucrose solutions (10–30%) and then frozen in OCT compound (Sakura Finetek). Sections were cut at a thickness of 4 μm with a CRYOSTAR NX70 instrument (ThermoFisher) and then immunostained. For Airyscan observation, a portion of each embryo including the node was excised and mounted in mounting medium with a refractive index of 1.52 (ProLong Glass, ThermoFisher) on a glass slide fitted with a silicone rubber spacer (thickness of 100 µm; 6-9085-12, ASONE) and was then covered with a coverslip for super-resolution imaging (No.1 S HT, Matsunami). The node was observed with a Zeiss LSM 880 with Airyscan confocal microscope equipped with 63× (Plan-APOCHROMAT 63× Oil for SR 1.4 N.A.) and 100× (alpha-Plan-APOCHROMAT 100 Oil for SR 1.46 N.A.) lenses. 3D images were recorded with a depth along the *z*-axis of 168 nm for 100× and 185 nm for 63×. After Airyscan processing, fluorescence intensity along the *z*-axis was measured for each pixel and the center of localization was determined by 1D Gaussian fitting. The distance along the *z*-axis between the centers of the red and green channels was calculated and averaged for the entire region of each cilium. Chromatic aberration was measured with the use of TetraSpeck fluorescent beads (diameter of 0.1 µm; T7279, ThermoFisher) and corrected for precisely. In images of frozen sections, chromatic aberration was corrected by the channel alignment function in Zen software (Carl Zeiss). 3D-STED imaging was performed with a TCS SP8 STED 3X instrument (Leica) equipped with a 100× lens (HC PL APO 100×/1.40 Oil STED white). Immunostaining was performed as described above but with secondary antibodies conjugated with STAR ORANGE (1:200 dilution, anti-chick; Abberior) or STAR RED (1: 200 dilution, anti-mouse; Abberior). Each embryo was also observed as described immediately above but with the use of TDE-DABCO mounting medium (97% thiodiethanol, 2.5% DABCO). The STED images were taken with pulsed 561 nm and 633 nm excitation laser lines and pulsed 775 nm depletion laser with 60% *z*-donut. The raw images were further processed by classical maximum likelihood estimation deconvolution algorithm to remove the haze from STED immunity fraction (Huygens, Scientific Volume Imaging). Angular distribution of the intensity in Pkd2 signal was analyzed with codes written in IgorPro 8 (WaveMetrics). The azimuth angle (θ) between each pixel, which has a green signal (Pkd2::Venus) above the threshold, and the center of red signals (acetylated tubulin) were calculated in each *z*-section. The green intensity is integrated over each 45° in each cilium. The integrated intensity is normalized as the sum of intensity to be one.

### FIB-SEM

Samples for FIB-SEM were prepared using Spurr’s Low-Viscosity resin according partially modified to previously described^39^ and observed. *Kif3a*^−/−^ embryos at the one-somite stage harboring an *NDE*-*Kif3a* transgene were collected in ice-cold PBS and fixed immediately by incubation overnight at 4°C in fresh 0.1 M cacodylate buffer (pH 7.4) containing 2% glutaraldehyde and 4% paraformaldehyde. They were then incubated in ice-cold 2% OsO_4_ for 2 h and embedded in resin. After smoothing of the surface of the embedded samples with an ultramicrotome, they were observed with a Helios G4 UC instrument (ThermoFisher). 3D reconstruction and segmentation were performed using Amira software Ver. 2020.2 (ThermoFisher).

### Mechanical modeling of ciliary membrane deformation

To measure the mechanical strain of the ciliary membrane, we first determined the center line of the same cilium with and without nodal flow. The radius of the cilium was assumed to be sufficiently small relative to its length for the Bernoulli-Euler hypothesis to hold. Material points of the ciliary membrane were then defined normal to the center line. The ciliary membrane was discretized with 2160 triangular elements, and, according to the thin shell theory^40^, the membrane strain was determined from the deformation in the presence of nodal flow, with that in the absence of the flow as the reference. A membrane material point is defined by two surface curvilinear coordinates (ξ^1^, ξ^2^) that correspond to position **X**(ξ^1^, ξ^2^) in the reference state and to position **x**(ξ^1^, ξ^2^) in the deformed state. The local covariant base vectors for both the reference and deformed states are then defined as tangential to lines of constant ξ^α^:

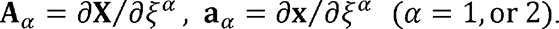

The associated contravariant base vectors **a**^α^ are also defined by 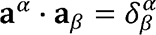 (likewise for **A**^α^), where 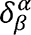 is the Kronecker tensor. The surface gradient of the transformation **F** is then given by

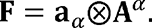

Local deformation of the membrane is estimated by the Green-Lagrange deformation tensor **e**

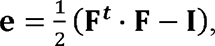

where **I** is the 2D identity tensor. The deformation invariants are then given by

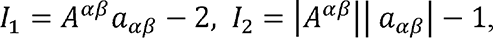

where 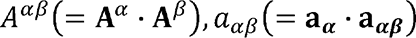 are the contravariant and covariant metric tensors in the reference and deformed states, respectively, and |*a_αβ_*| is the determinant of *a_αβ_*. The second invariant of the strain tensor was used as a representative quantity indicating membrane biaxial deformation. Assuming that the membrane is a 2D isotropic material, the Cauchy tension tensor is given by a strain energy function. We applied the Skalak model^26^ for the membrane constitutive law. This model represents large resistance to area change and large deformation in shear and is widely adopted for modeling of biomembranes. The two principal tensions τ_1_ and τ_2_ are given by

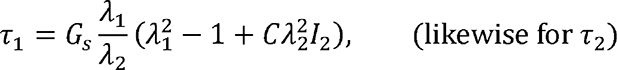

where λ_1_ and λ_2_ are the two principal in-plane stretch ratios, *G*_s_ (= 4 µN/m) (ref. ^41^) is the shear elastic modulus, and *C* (= 10^5^) (ref. ^42^) is a dimensionless material coefficient that measures the resistance to area dilation.

## Statistical analysis

Statistical analysis was performed and graphs were drawn with the use of IgorPro 8 (WaveMetrics). 3D reconstructed images were generated with the use of Imaris (Oxford Instruments). Statistical tests adopted are described in the figure legends. A *P* value of <0.05 was considered statistically significant.

## Data Availability

The data supporting the findings of this paper are available from the corresponding authors upon reasonable request.

## Code Availability

Code used in this study is available from the corresponding authors upon reasonable request.

## Acknowledgments

We thank Y. Kiyosue for support with microscopy systems; K. Kawaguchi and members of his laboratory as well as S. Nonaka for discussion; Laboratory for Ultrastructural Research (RIKEN BDR) for technical support; and K. Takaoka, T. Ide, and H. M. Takase for technical advice. This study was supported by grants from the Ministry of Education, Culture, Sports, Science, and Technology (MEXT) of Japan (no. 17H01435) and from Core Research for Evolutional Science and Technology (CREST) of the Japan Science and Technology Agency (JST) (no. JPMJCR13W5) to H.H.; by a Grant-in-Aid (no. 21K15096) from the Japan Society for the Promotion of Science (JSPS) to T.A.K; by RIKEN Special Postdoctoral Researcher Program to T.A.K; by a grant from Precursory Research for Embryonic Science and Technology (PRESTO) of the Japan Science and Technology Agency (no. JPMJPR2142) to T.O; and by RIKEN Cluster for Science, Technology and Innovation Hub (RCSTI). 3D-STED microscopy was supported by grants from JST (no. JPMJM2025-15, JPMJCR20E2, JPMJCR15G2, JPMJCR1851) and grants from JSPS (no. 19H05794, 16H06280) to Y.O.

## Author contributions

T.A.K. and T.N. designed experiments with optical tweezers. T.A.K. performed biophysical experiments with mouse embryos and analyzed the data. T.O. and T. Ishikawa are responsible for theoretical analysis of immotile cilia. Y.I. and H.N. generated transgenic mice. S.H. genotyped transgenic mice. K. Mizuno helped with analysis of Ca^2+^ transients and HILO imaging. K. Minegishi helped with analysis of *Dand5* mRNA degradation. E.K. examined ANE-LacZ activity. X.S. performed immunostaining for Pkd2::Venus. T. Itabashi and A.H.I. performed FIB-SEM analysis of immotile cilia. Y.O. assisted with STED analysis. T.A.K., T.O., and H.H. conceived the project and wrote the paper.

## Competing interests

The authors declare no competing interests.

## Additional information

**Supplementary information** The online version contains supplementary material available at https:

**Correspondence and requests for materials** should be addressed to Takanobu A. Katoh, Toshihiro Omori, or Hiroshi Hamada.

## Extended Data Figures

**Extended Data Fig. 1.**
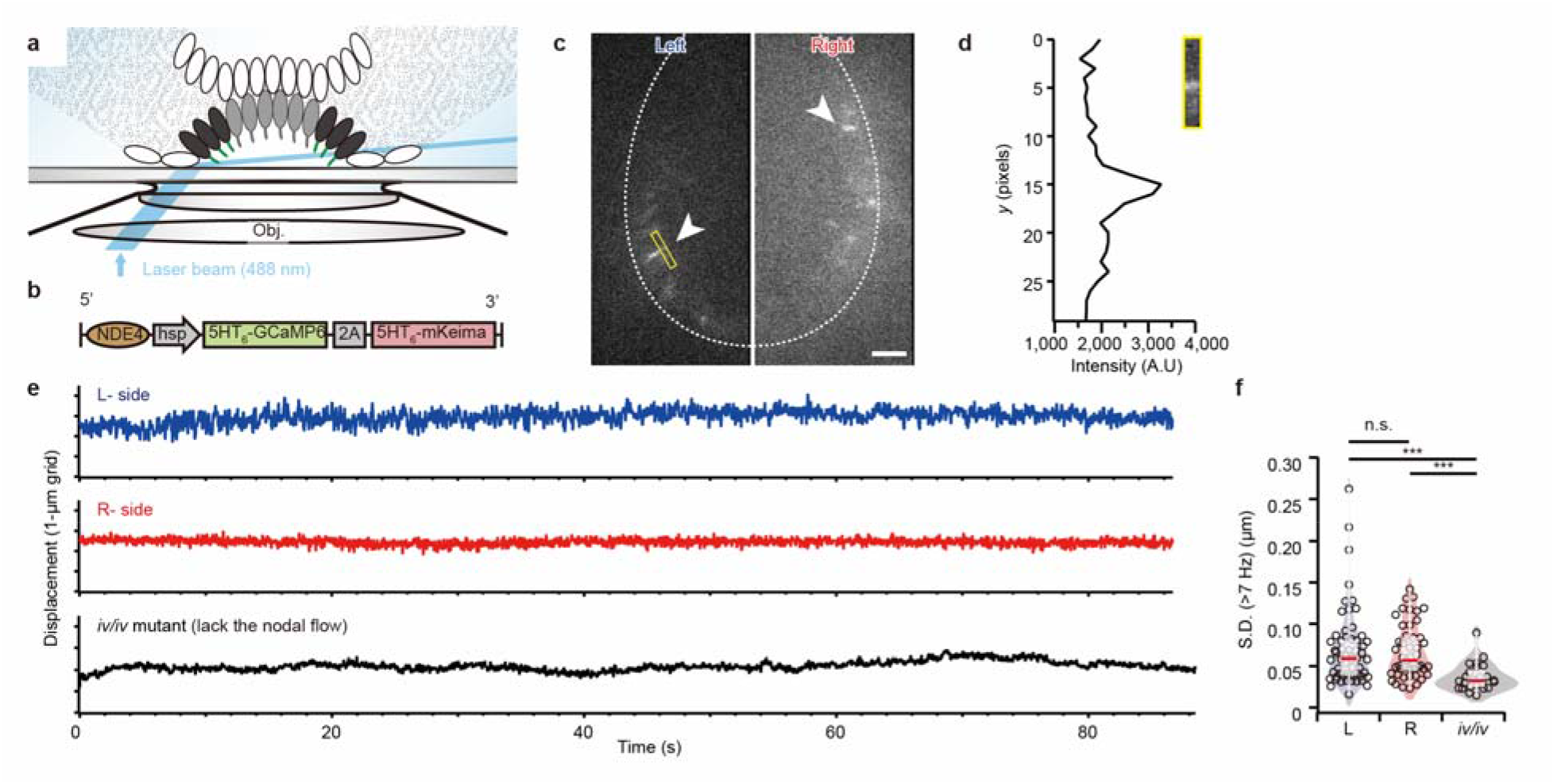
High-speed imaging of immotile cilia by HILO microscopy. **a**, Schematic of the experimental setup. The blue line represents the illumination beam for highly inclined and laminated optical sheet (HILO) microscopy. Immotile cilia at the mouse node expressing GCaMP6 was excited with a thin sheet of laser beam. Fluorescence signals were detected with an EM-CCD camera with a time resolution of 29.2 ms (34.3 frames/s). Obj., objective lens. **b,** Schematic of the transgene adopted for these experiments. These experiments were performed with embryos harboring the *NDE4-hsp-5HT_6_-GCaMP6-2A-5HT_6_-mKeima* or the *NDE4-hsp-5HT_6_-GCaMP6-2A-5HT_6_-mCherry* transgene. **c**, GCaMP6 fluorescence image obtained by HILO illumination of the node of a wild-type mouse embryo at the zero-somite stage harboring the transgene shown in **b** (see Supplementary Video 1). Scale bar, 10 µm. **d**, GCaMP6 intensity profile of a single immotile cilium. The center of intensity was tracked with sub-micrometer accuracy, and shown in **e**. A.U., arbitrary units. **e**, Trajectories of the tips of immotile cilia on the left and right sides of a wild-type embryo at the zero-somite stage as well as of a cilium of an *iv/iv* mutant embryo at the two-somite stage. **f**, Standard deviation (S.D.) of high-frequency (>7 Hz) components in the trajectories of immotile cilia, which reflects motion of the cilia, for wild-type (L, 66 ± 44 nm; R, 65 ± 33 nm) and *iv/iv* (29 ± 15 nm) embryos (*n* = 57, 47, and 24 cilia, respectively). Data are means ± s.d. \*\*\**P* < 0.001, n.s. (Mann-Whitney *U* test).

**Extended Data Fig. 2.**
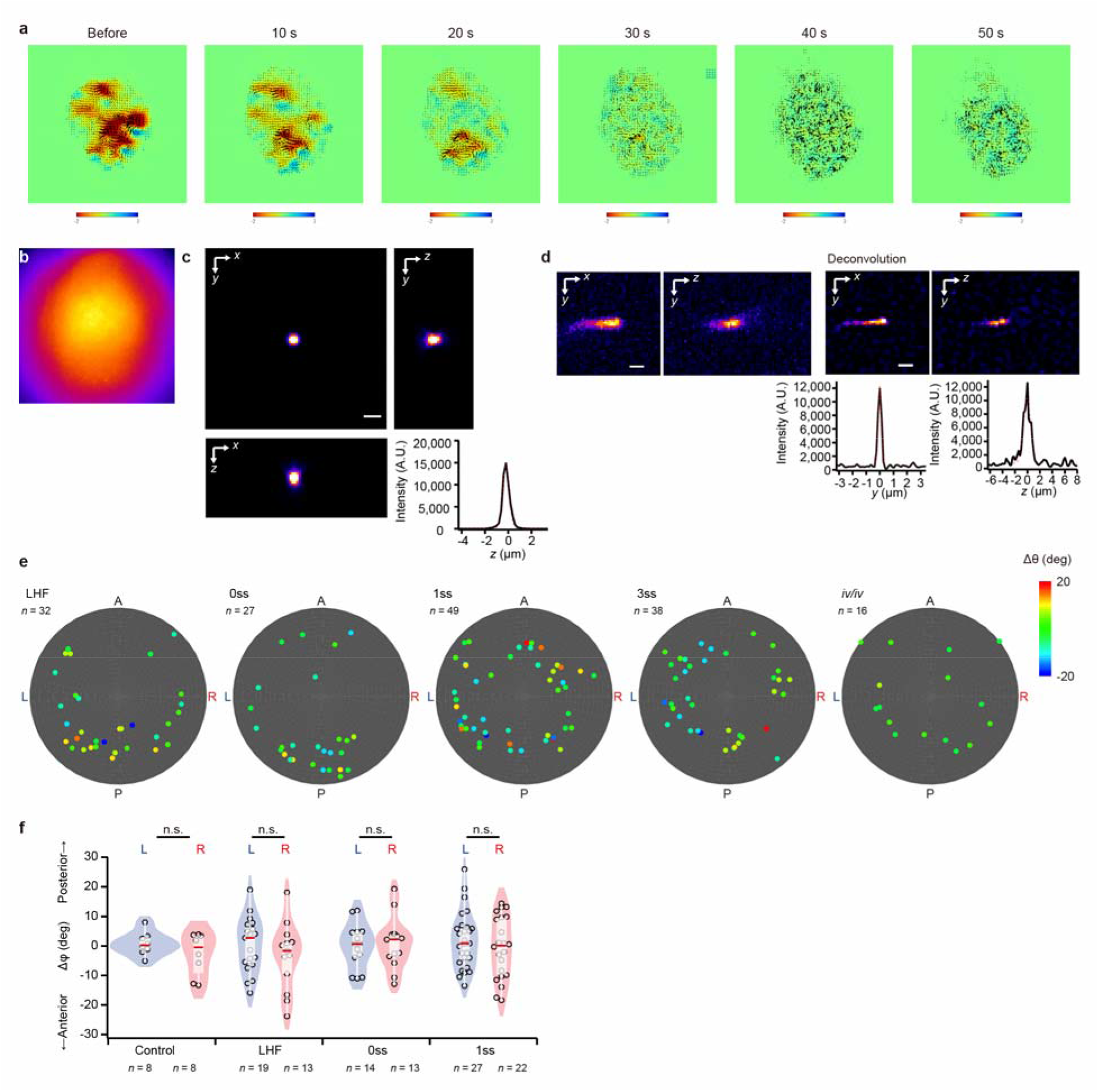
Bending of immotile cilia in response to nodal flow. **a**, PIV analysis of nodal flow in a wild-type embryo at the two-somite stage both before and after UV irradiation for the indicated times. Nodal flow was completely lost after UV irradiation for ∼40 s. **b**, The area of UV irradiation was confirmed with an embryo embedded in medium containing a high concentration of fluorescent beads. Compared with the entire node area shown in a background bright-field image, the UV irradiation area covers the central region of the node where pit cells with motile cilia are located. **c**, Point spread function for the deconvolution calculation. Scale bar, 10 µm. **d**, Images before and after deconvolution for the left-side cilium shown in Figure 1f. Intensity profiles of the cilium along the *y-*and *z-*axes are also shown. Full width at half maximum (FWHM) values along the *z*-and *y*-axes are 1.72 and 0.46 µm, respectively, after deconvolution. **e**, Δθ and the location within the node are summarized for all immotile cilia examined for wild-type embryos at the LHF as well as zero-, one-, and three-somite stages as well as for *iv/iv* embryos at the LHF to the two-somite stage. **f**, Change in the azimuth angle (Δφ), which represents bending of an immotile cilium along the A-P axis (see Fig. 1g), between before and after UV irradiation, for the indicated embryos. Data for *iv/iv* embryos at the LHF to two-somite stage are shown as a control. n.s., Mann-Whitney *U* test.

**Extended Data Fig. 3.**
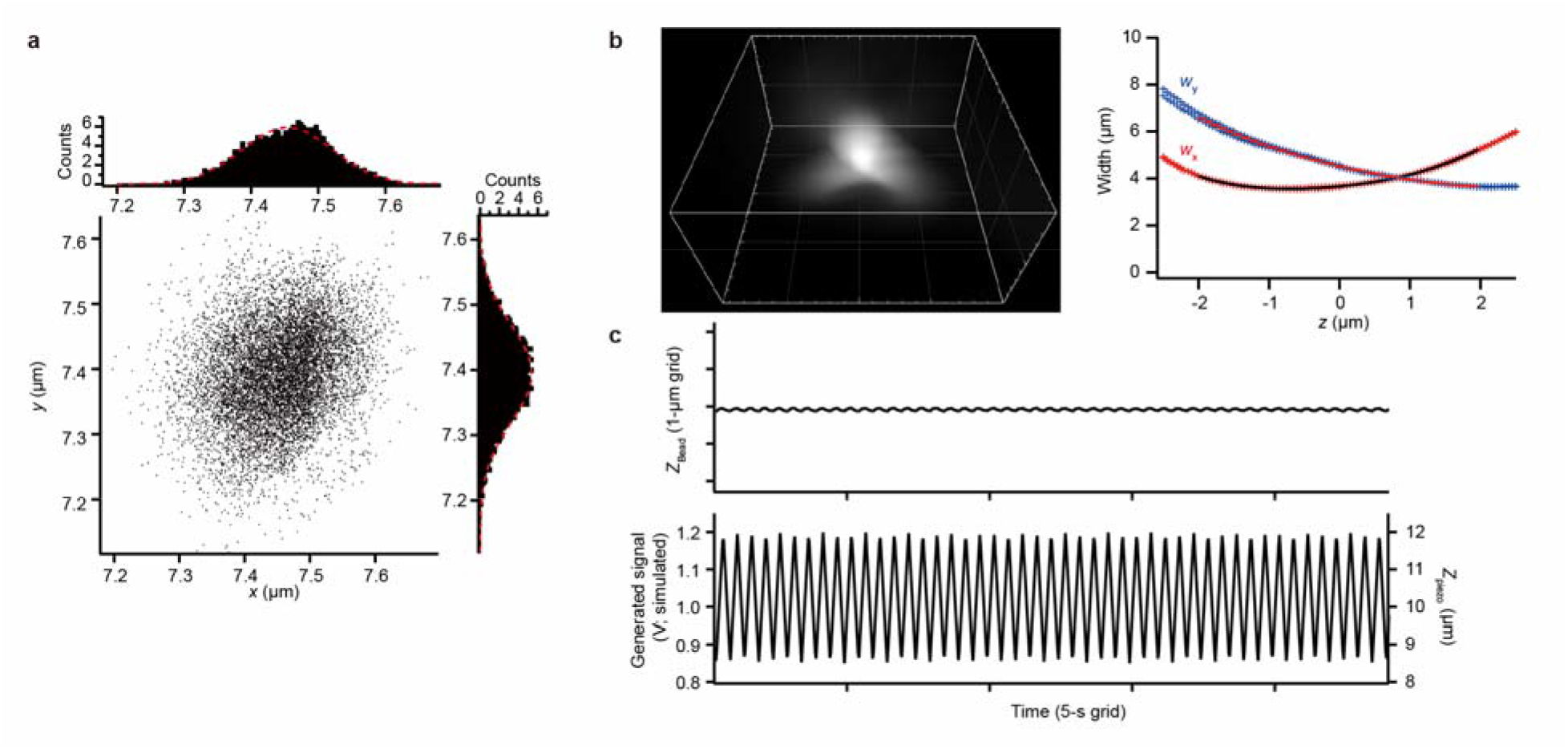
Calibration of optical tweezers and 3D tracking microscopy with 3.5-µm beads. **a**, Trajectories of 3.5-µm-diameter beads trapped by a laser power of 11.8 mW, and the probability of bead localization in the *x* and *y* directions. The center of each bead was estimated from a bright-field image. The red dotted line indicates Gaussian fitting for calculation of trapping stiffness, which was estimated as 70 ± 12 and 51 ± 10 pN µm^−1^ W^−1^ along the *x* and *y* directions, respectively (*n* = 5 determinations). Maximal trapping force imposed by a laser power of 800 mW was estimated from the product of the spring constant and the range of trapping as ∼±12 pN in the *xy* plane (*n* = 5), which is sufficient to bend a cilium. **b**, Point spread function of fluorescent beads (diameter of 4 µm) observed by 3D single-particle tracking microscopy (left panel), and the resultant calibration curve (right panel). Grid size, 5 µm. **c**, The *z*-motion of beads (diameter of 4 µm) trapped by a laser power of 800 mW as observed by 3D single-particle tracking microscopy (upper panel), and the change in voltage applied to the piezo *z*-stage (lower panel). The small displacement of the trapped bead in the *z* direction confirms that the displacement between the center of the bead and the objective was constant even though the piezo *z*-stage was moving a distance of ±1.75 µm.

**Extended Data Fig. 4.**
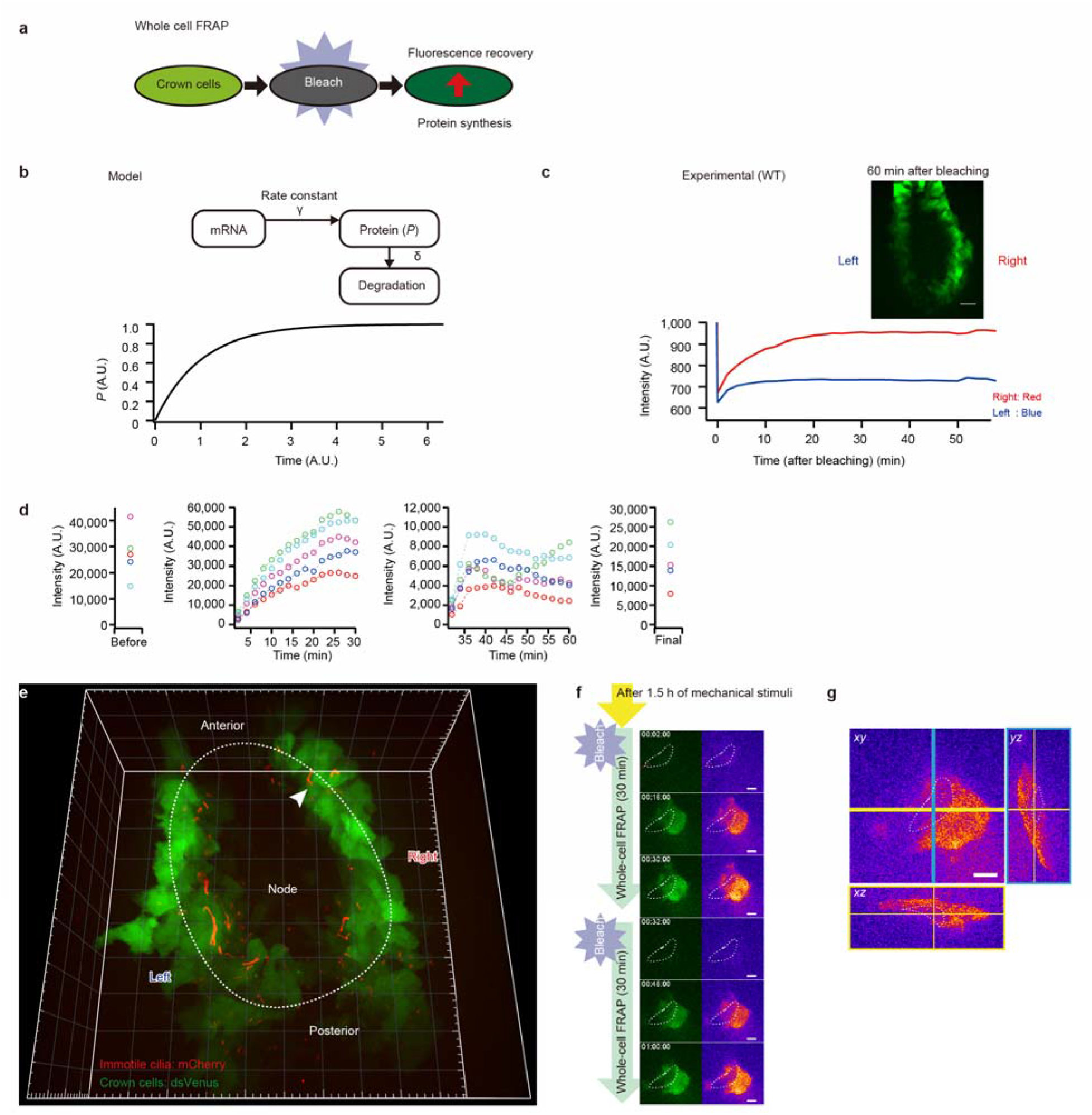
Whole-cell FRAP measurement. **a**, Schematic of whole-cell FRAP measurement. Recovery of fluorescence after the entire region of a cell is subjected to photobleaching depends on protein synthesis. **b**, A simple model in which mRNA is translated to yield the fluorescent protein with a rate constant γ, and the protein is degraded with a rate constant δ. This model is consistent with the actual fluorescence recovery curves for the experiment shown in **c**. *P* represents the amount of protein (see methods). **c**, The node region of a wild-type (WT) embryo harboring the *NDE4-hsp-dsVenus-Dand5-3′-UTR* transgene at the three-somite stage shows a more pronounced recovery of fluorescence on the right side than on the left side. Scale bar, 20 μm. The fluorescence recovery reaches a plateau at ∼20 min on both sides. **d**, Actual fluorescence intensity of cells analyzed in Figure 2f. Whereas the recovery of fluorescence is normalized in Figure 2f, the intensity values here do not account for differences in acquisition settings such as exposure time. **e**, 3D image of the node of a wild-type embryo harboring the *NDE4-hsp-dsVenus-Dand5-3′-UTR* transgene. The white arrowhead indicates the cell with the cilium stimulated in **f**. Grid size, 20 μm. **f**, Fluorescence recovery on the right side of the embryo in **e**. The cell with the stimulated cilium located on the right side of the node (indicated by the white dotted line) shows weak recovery of fluorescence. 2D sections from 3D images are shown (see Supplementary Video 7). Scale bars, 10 μm. **g**, Sections (*xy*, *xz*, and *yz*) of the 3D image of the embryo in **f** acquired at 1 h after stimulation. Scale bar, 10 μm.

**Extended Data Fig. 5.**
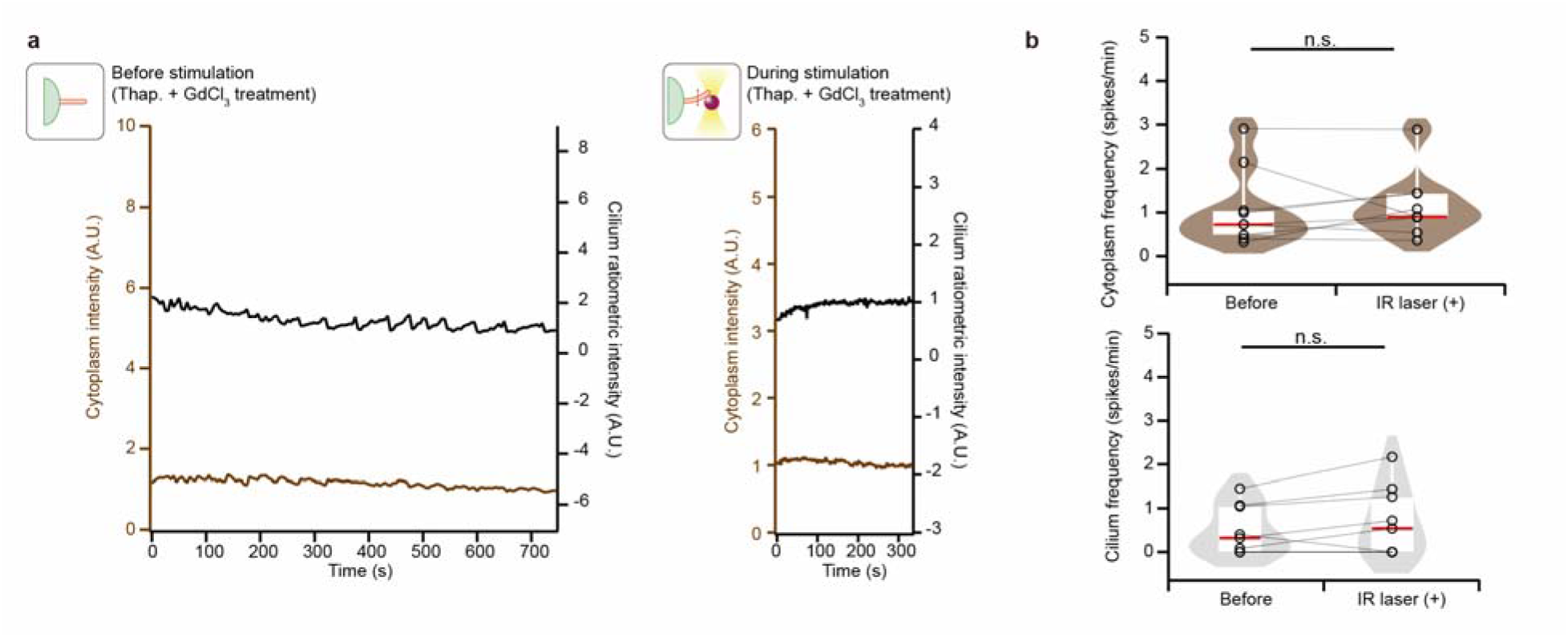
Optical tweezers did not affect intracellular Ca^2+^ dynamics. **a**, GCaMP6 fluorescence intensity in the cytoplasm and the GCaMP6/mCherry fluorescence intensity ratio in the immotile cilium of a crown cell in a wild-type embryo exposed to thapsigargin (Thap.) and GdCl_3_ before and during mechanical stimulation of the cilium. In the presence of thapsigargin and GdCl_3_, Ca^2+^ transients in the cilium and cytoplasm are lost, as described previously^21^. The fluorescence intensity remains almost constant during stimulation, which ensures that Ca^2+^ measurement during stimulation is valid and motion artifacts can be ignored. **b**, Infrared laser irradiation associated with the use of optical tweezers did not affect Ca^2+^ transients in the cytoplasm and cilia. Frequencies of ciliary and cytoplasmic Ca^2+^ transients were measured on the basis of GCaMP6/mCherry fluorescence intensity ratio and GCaMP6 fluorescence both before and during infrared laser irradiation of the distal portion of an immotile cilium (*n* = 9 cells and cilia from nine embryos). n.s., Wilcoxon signed-rank test.

**Extended Data Fig. 6.**
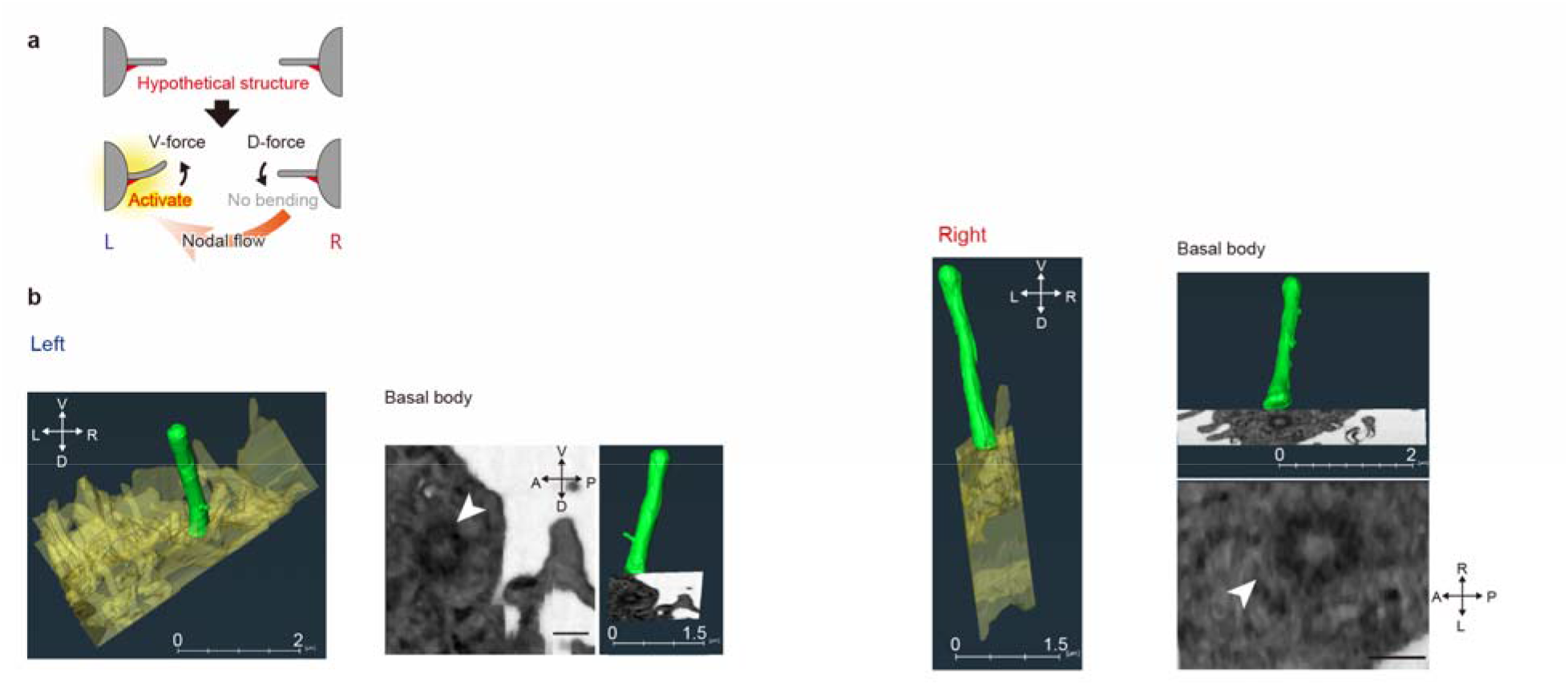
FIB-SEM observation of immotile cilia at the node. **a**, Hypothetical model to explain why immotile cilia on the left side of the node preferentially bend toward the ventral side. A structure (such as a basal foot) that is asymmetrically localized along the D-V axis of the cilium might promote preferential bending toward the ventral side in response to the leftward nodal flow. **b**, FIB-SEM images of immotile cilia in *Kif3a*^−/−^ embryos at the one-somite stage harboring an *NDE*-*Kif3a* transgene, which possess immotile cilia only at the node^2^. Immotile cilia on the left and right sides of the node are shown. Green and yellow regions in the 3D reconstructed images represent the axoneme and surface of the cell body, respectively. A cross-section at the base of each cilium shows the basal body (white arrowhead). An asymmetric structure at or near the base of immotile cilia was not detected. Scale bars, 0.25 µm for cross-section image of the basal body.

**Extended Data Fig. 7.**
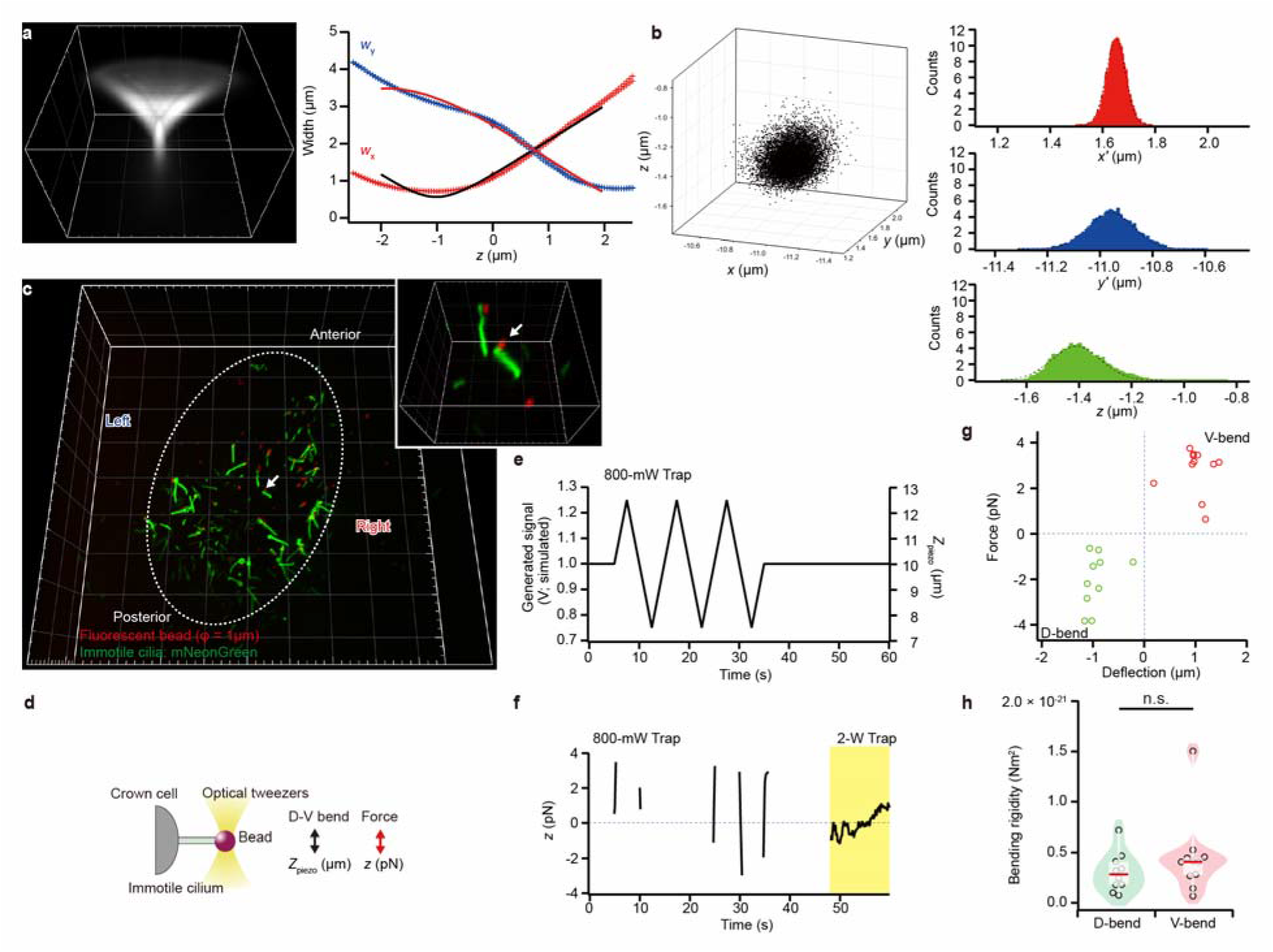
Flexural rigidity of immotile cilia at the node. **a**, Point spread function of beads with a diameter of 1 µm observed by 3D single-particle tracking microscopy (left panel), and the resultant calibration curve (right panel). **b**, 3D trajectories of beads with a diameter of 1 µm trapped by a laser power of 30.48 mW (left panel), and the probability of localization of the beads in *x’*, *y’*, and *z* directions (right panel). We defined *x’*-and *y’*-axes to correct for anisotropy of trapping stiffness in the *xy* plane that results from aberration in the system. The black dotted lines indicate Gaussian fitting for calculation of trapping stiffness. Trapping stiffness was estimated as 97 ± 9, 19 ± 1, and 16 ± 1 pN µm^−1^ W^−1^ along the *x’*, *y’*, and *z* directions, respectively (means ± s.d., *n* = 5 determinations). **c**, Attachment of fluorescent beads (diameter of 1 µm) to immotile cilia at the node. mNeonGreen (green) is expressed in the cilia of a wild-type mouse embryo, and a red fluorescent bead is attached to the tip of a cilium (arrow). Grid size, 20 and 5 µm for the main and magnified panels respectively. **d**, Schematic of the experimental protocol for **e** to **h**. A bead attached to a cilium was trapped by optical tweezers and manipulated to induce D-V bending of the cilium. The resistance force of the cilium along the *z*-axis was measured. **e**, Change in voltage applied to the piezo *z*-stage in D-V bending experiments. **f**, Resistance force of cilia along the *z*-axis for D-V bending. Each cilium was subjected to bending of ±2.5 µm along the *z*-axis with a 0.1-Hz ramp wave. Force along the *x*-, *y*-, and *z*-axes was calculated from the 3D displacement of the bead from the center of the trapping position, which was determined by 3D single-particle tracking microscopy. **g**, Relationship between the deflection of cilia and the applied load by optical tweezers. The positive and the negative direction of each axis indicate the ventral and the dorsal direction, respectively. Red and green circles represent results for the ventral and dorsal bending, respectively. **h,** Bending rigidity in the ventral and the dorsal direction. Bending rigidity of immotile cilium was estimated as 4.5 ± 3.8 × 10^-22^ and 3.0 ± 1.9 × 10^-22^ Nm^2^ for dorsal and ventral bending (*n* = 11 and 10 determinations from three cilia). n.s., Mann-Whitney *U* test.

**Extended Data Fig. 8.**
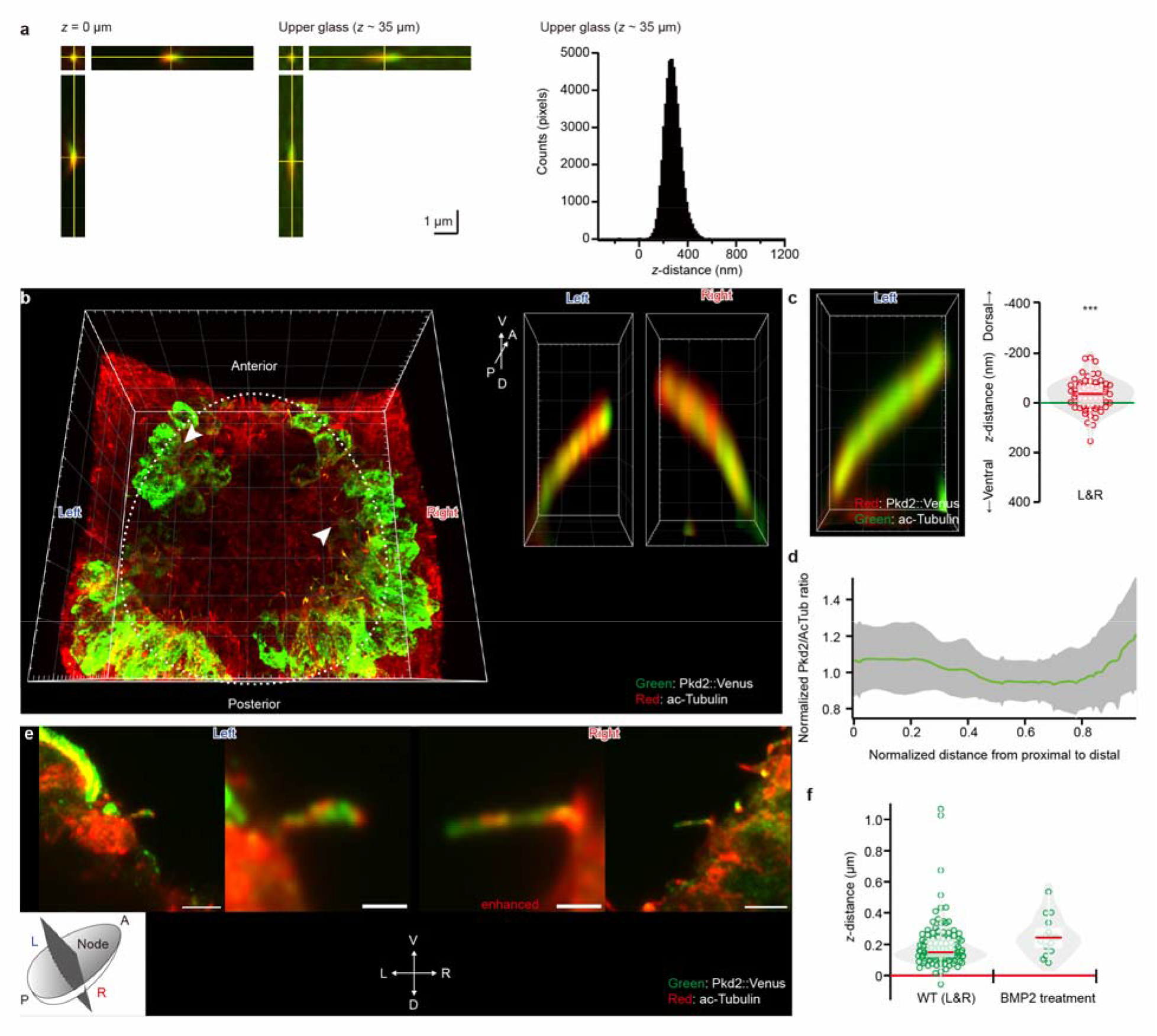
Measurement of the polarized localization of the Pkd2 channel in immotile cilia. **a**, The two-color 3D point spread function of the Airyscan microscope was measured at the glass surface (*z* = 0 µm) and *z*-axis distance of ∼35 µm (left panels). For quantification of chromatic aberration along the *z*-axis, the distance along this axis between red and green channels for each pixel was measured by Gaussian fitting (right panel) and determined to be 272 ± 78 nm (mean ± s.d., *n* = 50917 pixels from 5 determinations). **b**, A wild-type mouse embryo harboring an *NDE2-hsp-Pkd2-Venus* transgene^2^ was subjected to immunofluorescence analysis for detection of the Pkd2::Venus fusion protein and acetylated (ac)–tubulin at the node with an Airyscan microscope equipped with a 63× objective (main image). Magnified views of D-V sections with a 100× objective shown on the right suggested a preferential localization of Pkd2::Venus at the dorsal side of cilia on both the left and right sides of the node. Grid size, 10 and 1 µm for the main and magnified panels, respectively. **c**, Immunofluorescence image of Pkd2::Venus (red) and acetylated tubulin (green) at the node of a wild-type embryo harboring the *NDE2-hsp-Pkd2-Venus* transgene as obtained by Airyscan microscope with a 100× objective (left panel). The distance along the *z*-axis between the intensity centers of green and red fluorescence in longitudinal optical section of immotile cilia was measured by Gaussian fitting, after correction for chromatic aberration (right panel). The intensity center of Pkd2::Venus (red) was significantly polarized toward the dorsal side of cilia on the left or right sides of the node. Such polarization was thus also detected when the fluorescent labels of the two secondary antibodies were switched relative to those used in Figure 4c (*n* = 61 cilia from six embryos). Grid size, 1 µm. \*\*\**P* < 0.001 (One-sample *t*-test) **d**, Distribution of Pkd2 protein along the proximodistal axis of immotile cilia imaged as in **b**. The ratio of the immunofluorescence intensity of Pkd2::Venus to that of acetylated tubulin (AcTub) was calculated along the proximodistal axis of each cilium. After normalization of the length of the cilia and ratiometric intensity, data from 82 cilia were averaged, with the gray area indicating the standard deviation. **e**, Longitudinal sections of immotile cilia on the left and right sides of the node. Middle left and middle right panels show the magnified views of the left and right side cilium, respectively. Immunofluorescence images of Pkd2::Venus (green) and acetylated tubulin (red) obtained by Airyscan microscope with a 100× objective are shown. Pkd2 is localized to the dorsal side of each cilium. Scale bars, 3 µm (left and right panels) and 1 µm (middle left and middle right panels). **f**, The distance along the *z*-axis between the localization centers of Pkd2::Venus (green) and acetylated tubulin (red) was determined for immotile cilia on the left and right sides of the node of wild-type embryos incubated in the absence or presence of BMP2 from early bud stage to three-somite stage as in **b**. Preferential localization of Pkd2 at the dorsal side was maintained in the BMP2-treated embryos (*n* = 15 cilia from three BMP2-treated embryos and 107 cilia from 13 wildtype embryos).

**Extended Data Fig. 9.**
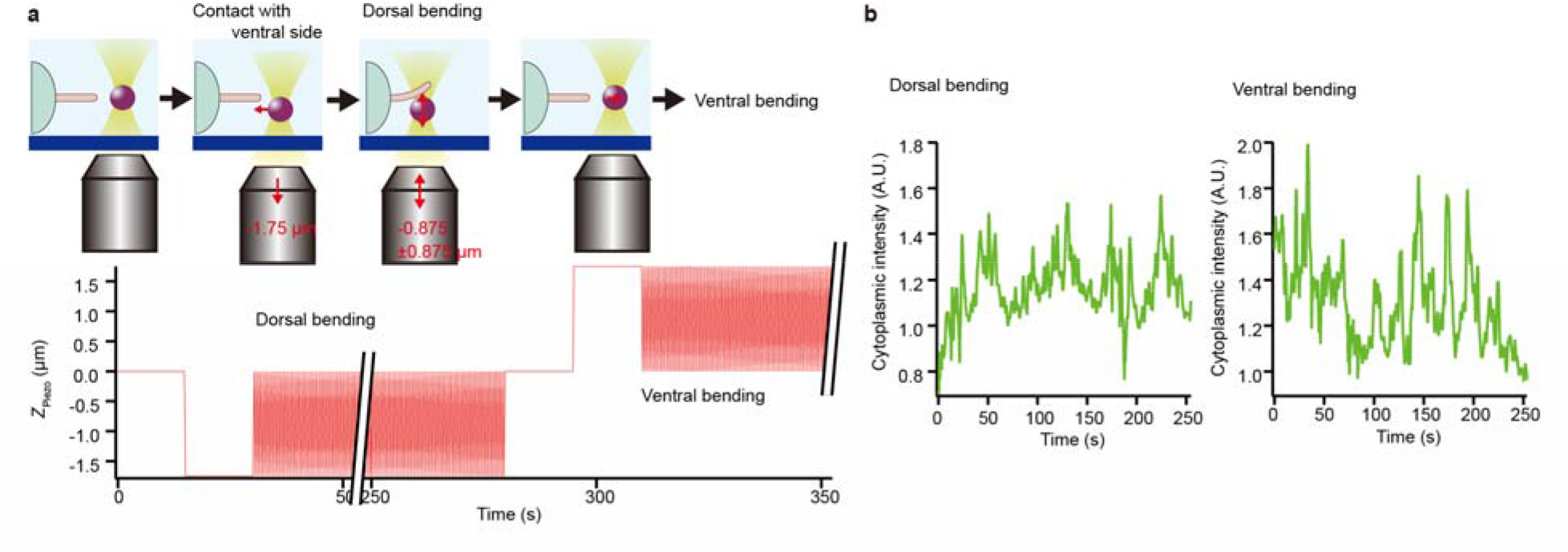
Ventral bending of immotile cilium preferentially leads to a Ca^2+^ response in cytoplasm. **a**, Motion of the piezo *z*-stage and experimental procedure. To apply dorsal bending to an immotile cilium at the node of an *iv/iv* embryo, a bead (diameter of 3.5 µm) was trapped and displaced to *z* = −1.75 µm (the same as a radius of the bead) and was contacted with the ventral side of the cilium (upper panel). The cilium was subjected to dorsal bending of –0.875 µm at 2 Hz with an amplitude of ± 0.875 µm for ∼250 s (cilium was bent 0-1.75 µm toward the dorsal side) and then to ventral bending of +0.875 µm at 2 Hz with an amplitude of ± 0.875 µm for ∼250 s (lower panel). **b**, Time course of the cytoplasmic intensity of Ca^2+^ imaging during dorsal and ventral bending of a cilium as in **a**. The frequency of Ca^2+^ transients was measured and is shown in Figure 4e (see also Supplementary Video 11).

**Extended Data Fig. 10.**
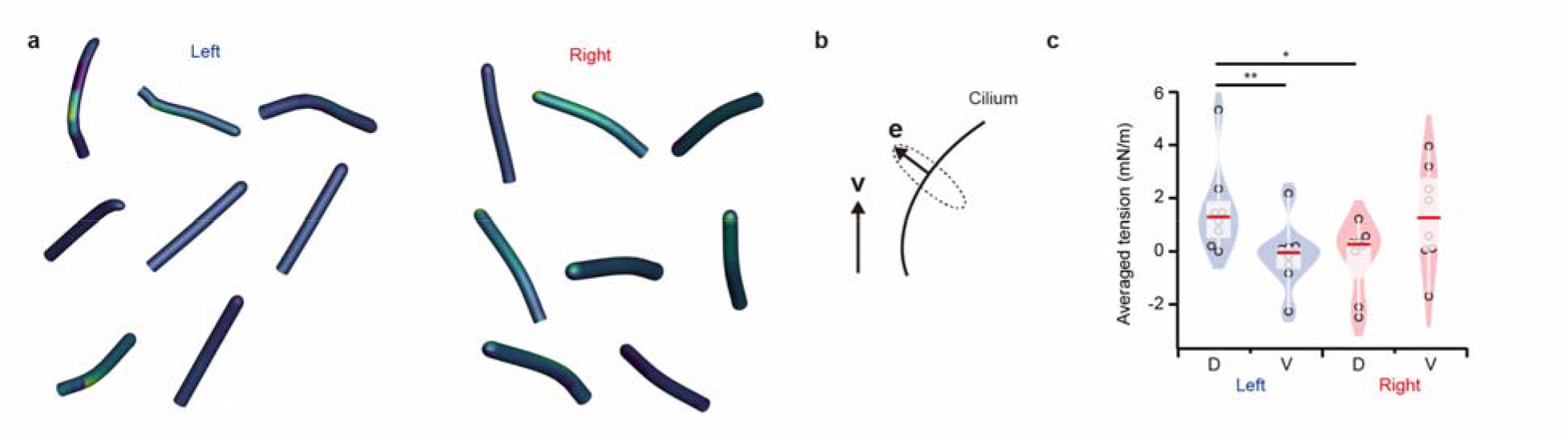
Modeling of an immotile cilium and estimation of membrane tension. **a**, Distribution of biaxial tension acting on the ciliary membrane (16 cilia from eight embryos). For mechanical modeling, the membrane was assumed to be a 2D isotropic hyperelastic material (see Methods). All immotile cilia on the left side show an increment of membrane tension on the dorsal side, where the Pkd2 channel is localized, whereas those on the right side rarely show such an increment. **b**, Determination of the dorsal and ventral sides of a cilium. The dorsal side of a cilium is determined by **e**· **v** < 0, where **e** is the unit normal vector from the center line to the membrane surface, and **v** is the unit vector along the *z*-axis (that is, the basis vector for the ventral direction). After determination of the dorsal and ventral regions, the strain and tension applied to the dorsal and ventral sides of the left and right cilia were compared by integrating and averaging in each region. **c**, Averaged tension on the dorsal and ventral sides of cilia. Averaged tension on the dorsal and ventral sides of each left-side cilium was estimated to be 1.6 ± 1.6 and –0.14 ± 1.2 mN/m (means ± s.d., *n* = 8), respectively, whereas the corresponding values for each right-side cilium were –0.24 ± 1.2 and 1.3 ± 1.8 mN/m (*n* = 8). \**P* < 0.05, \*\**P* < 0.01 (Student’s paired *t*-test).

## Notes

### Competing Interest Statement

The authors have declared no competing interest.

