## Supplementary information for "Immotile cilia of the mouse node sense a fluid flow–induced mechanical force for left-right symmetry breaking"

[7]Present address: Department of Cell Biology and Biochemistry, Division of Medicine, Faculty of Medical Sciences, University of Fukui, Eiheiji-cho, Yoshida-gun, Fukui, Japan

[8]Present address: Department of Molecular Therapy, National Institutes of Neuroscience, National Center of Neurology and Psychiatry, Kodaira, Tokyo, Japan.

**Supplementary Video 1. HILO imaging of immotile cilia at the node of a mouse embryo.** Images of GCaMP6 expressed in immotile cilia at the node of a wild-type mouse embryo at the zero-somite stage were obtained by HILO microscopy at a time resolution of 29.2 ms. Some cilia show an abrupt change in bending as a result of nodal flow. Scale bar, 10 μm.

**Supplementary Video 2. Loss of the beating of motile cilia at the node induced by UV irradiation.** Bright-field images of motile cilia at the node of a wild-type mouse embryo at the two-somite stage were obtained during UV irradiation. Irradiation starts at time “00:00” and results in a gradual decrease in beating frequency and an eventual complete loss of beating movement. Scale bar, 5 μm.

**Supplementary Video 3. Three-dimensional images of immotile cilia at the node before and after UV irradiation.** 3D images were obtained before and after UV irradiation of a wild-type mouse embryo at the two-somite stage. After 3D deconvolution, the edge of each cilium was detected and the change in the angle of the cilium was measured by ellipsoidal fitting.

**Supplementary Video 4. Three-dimensional imaging of the node before application of mechanical stimuli to immotile cilia.** The node of an *iv/iv* mouse embryo harboring the *NDE4-hsp-dsVenus-Dand5-3′-UTR* transgene, and therefore expressing mCherry in immotile cilia and dsVenus in the cytoplasm, is shown. The cilia subjected to mechanical stimulation are indicated.

**Supplementary Video 5. Mechanical stimulation of immotile cilia by optical tweezers.** Immotile cilia of crown cells of an *iv/iv* mouse embryo harboring the *NDE4-hsp-dsVenus-Dand5-3′-UTR* transgene were subjected to mechanical stimulation for 1.5 h with a bead that was trapped by optical tweezers and oscillated along the *z*-axis at 2 Hz with an amplitude of ±1.75 μm. Scale bar, 10 μm.

**Supplementary Video 6. Three-dimensional time-lapse imaging during whole-cell FRAP analysis of an *iv/iv* embryo.** Mechanical stimuli were administered to cilia of two crown cells for 1.5 h, after which all cells including the stimulated cells were subjected to two consecutive sessions of photobleaching and monitoring of the recovery of dsVenus fluorescence over 30 min.

The recovery of fluorescence in two crown cells whose immotile cilium were stimulated (white dotted line) was markedly less pronounced than was that in unstimulated cells.

**Supplementary Video 7. Time-lapse imaging during whole-cell FRAP analysis of a wild-type embryo.** Mechanical stimuli were administered to an immotile cilium of a crown cell on the right side of the node for 1.5 h, after which all cells including the stimulated cell were subjected to two consecutive sessions of photobleaching and monitoring of the recovery of dsVenus fluorescence over 30 min. The recovery of fluorescence in the crown cell whose immotile cilium was stimulated was markedly less pronounced than was that in unstimulated cells. Pseudo-colors of the right panel indicate the intensity of dsVenus fluorescence. Scale bar, 10 μm.

**Supplementary Video 8. Cytoplasmic $Ca^{2+}$ response to mechanical stimuli applied to the immotile cilium of a crown cell in an *iv/iv* embryo.** Cytoplasmic $Ca^{2+}$ transients were monitored with GCaMP6. Flow-independent $Ca^{2+}$ transients were observed in both the cytoplasm and cilium. The administration of mechanical stimuli to the cilium (arrowhead) resulted in an increase in the frequency of cytoplasmic $Ca^{2+}$ transients. Scale bar, 20 μm.

**Supplementary Video 9. Three-dimensional imaging of Pkd2 distribution in immotile cilia obtained by STED microscopy.** Immunofluorescence imaging of Pkd2::Venus and acetylated tubulin in a wild-type embryo was performed by STED microscopy. The Pkd2::Venus protein is preferentially localized at the dorsal side (the side facing the midline of the embryo) of immotile cilia on both left and right sides of the node.

**Supplementary Video 10. Three-dimensional imaging of Pkd2 distribution in immotile cilia obtained by confocal microscope with Airyscan detector.** Immunofluorescence imaging of Pkd2::Venus and acetylated tubulin in a wild-type embryo was performed by confocal microscope with Airyscan detector. The Pkd2::Venus protein is preferentially localized at the dorsal side of immotile cilia on both left and right sides of the node.

**Supplementary Video 11. $Ca^{2+}$ response to dorsal and ventral bending of immotile cilia.** The left panel is a raw movie obtained with an exposure of 200 ms. The right panel shows the running average with a five-frame window, which was used for the analysis shown in Figure 4e. The cilium was subjected to dorsal bending followed by ventral bending. Scale bar, 10 µm.
